# Multiplexed Non-invasive *in vivo* Imaging to Assess Metabolism and Receptor Engagement in Tumor Xenografts

**DOI:** 10.1101/758383

**Authors:** Alena Rudkouskaya, Nattawut Sinsuebphon, Marien Ochoa, Joe E. Mazurkiewicz, Xavier Intes, Margarida Barroso

**Affiliations:** Department of Cellular and Molecular Physiology, Albany Medical College, Albany, New York 12208, USA; Department of Biomedical Engineering, Rensselaer Polytechnic Institute, Troy, New York 12180, USA; Department of Neuroscience and Experimental Therapeutics, Albany Medical College, Albany, NY 12208, USA

**Keywords:** Targeted drug delivery, preclinical imaging, optical imaging, multiplexed imaging, metabolic imaging, breast cancer, lifetime imaging, FRET

## Abstract

Following an ever-increased focus on personalized medicine, there is a continuing need to develop preclinical molecular imaging modalities to guide the development and optimization of targeted therapies. To date, non-invasive quantitative imaging modalities that can comprehensively assess simultaneous cellular drug delivery efficacy and therapeutic response are lacking. In this regard, Near-Infrared (NIR) Macroscopic Fluorescence Lifetime Förster Resonance Energy Transfer (MFLI-FRET) imaging offers a unique method to robustly quantify receptor-ligand engagement *in vivo* and subsequent intracellular internalization, which is critical to assess the delivery efficacy of targeted therapeutics. However, implementation of multiplexing optical imaging with FRET *in vivo* is challenging to achieve due to spectral crowding and cross-contamination. Herein, we report on a strategy that relies on a dark quencher that enables simultaneous assessment of receptor-ligand engagement and tumor metabolism in intact live mice. First, we establish that IRDye QC-1 (QC-1) is an effective NIR dark acceptor for the FRET-induced quenching of donor Alexa Fluor 700 (AF700) using *in vitro* NIR FLI microscopy and *in vivo* wide-field MFLI imaging. Second, we report on simultaneous *in vivo* imaging of the metabolic probe IRDye 800CW 2-deoxyglucose (2-DG) and MFLI-FRET imaging of NIR-labeled transferrin FRET pair (Tf-AF700/Tf-QC-1) uptake in tumors. Such multiplexed imaging revealed an inverse relationship between 2-DG uptake and Tf intracellular delivery, suggesting that 2-DG signal may predict the efficacy of intracellular targeted delivery. Overall, our methodology enables for the first time simultaneous non-invasive monitoring of intracellular drug delivery and metabolic response in preclinical studies.

## Introduction

Skyrocketing cost of new targeted drug development combined with high failure rate of clinical trials call for a new paradigm of drug delivery assessment in living intact small animals. Especially, during the preclinical stage, it is critical to characterize the extent and duration of drug-target engagement in an undisturbed tumor environment to reflect on potential clinical efficacy.^1–3^ However, a major challenge in preclinical molecular imaging is the lack of multiplexing approaches that can directly assess intracellular anti-cancer drug delivery while also reporting on drug efficacy via tumor metabolic signatures in live animal models non-invasively and longitudinally.^4–7^ Indeed, current standard approaches are either destructive or incapable to discriminate between cell-associated drugs and unbound drugs residing in the extracellular space.^8–11^

Monitoring drug biodistribution and quantification in preclinical *in vivo* studies greatly benefits from Fluorescence Lifetime Imaging (FLI). FLI is widely regarded as the most robust means to utilize Förster Resonance Energy Transfer (FRET) to study receptor-ligand target engagement in living systems. Upon donor excitation, FLI estimates FRET occurrence by determining the reduction of the fluorescence lifetime of the donor molecule, when in nanometer (2-10nm) proximity of an acceptor molecule.^12–14^ When applied to receptor-ligand systems, FRET occurs when donor-labeled and acceptor-labeled ligands/antibodies bind to dimerized or cross-linked receptors.^15–21^ Hence, FLI FRET acts as a direct reporter of receptor engagement and internalization via the measurement of the fraction of labeled-donor entity undergoing binding to its respective receptor and subsequent internalization.^18, 21^ Recently, our group has pioneered Near-Infrared (NIR) Macroscopic FLI-FRET (MFLI-FRET) to quantitatively monitor ligand-target engagement, a critical component of targeted drug delivery, in live and intact animals.^22–24^ Using the transferrin (Tf)-transferrin receptor (TfR) system as a biological model, we demonstrated that MFLI-FRET quantitatively reported on intracellular delivery of the drug carrier ligand, providing a non-invasive direct readout of the true payload delivered to pathological cells in live intact animals.^22^ Hence, MFLI-FRET is poised to become a mainstream analytical tool to assess target engagement of the ligand- or antibody to their respective receptors in preclinical studies. However, to extend the utility of this methodology to the monitoring of multiple targets simultaneously it is crucial to enable the quantitative imaging of multiple fluorescent reporters simultaneously in the same model system. For instance, to accurately quantify the tumor-targeted drug delivery and efficacy in preclinical research, there is a pressing need to develop a protocol to simultaneously assess tumor metabolism in response to drug-target engagement across the undisturbed whole tumor in live intact animals. This can be achieved with the metabolic probe IRDye 800CW 2-deoxy glucose (2-DG),^25^ which is a fluorescent version of the ubiquitously used in clinical PET imaging ^18^F-fluorodeoxyglucose (FDG).^26^ However, the spectral crowding associated with the acceptor fluorophore required in MFLI-FRET limits the potential to perform multiplexing without crosstalk and FRET signal contamination. Indeed, for optimal FRET pair performance, the donor emission and acceptor excitation spectra need to have a large spectral overlap along with enough separation to avoid signal crosstalk. These requirements render multiplexing FRET with other fluorescence intensity imaging particularly difficult to achieve. A possibility would be to use FRET pairs and other fluorophores that are widely spectrally separated such as NIR and CFP-YFP biosensors.^27^ Alternatively, lifetime multiplexing could be achieved by using fluorophores displaying similar excitation wavelengths but distinctly separated lifetimes.^28^ Other studies have multiplexed FRET in single cells,^29–31^ as well as in an ear edema model *in vivo*^32^ using systemic administration of the fluorescent probes. However, for the NIR region, which is most suitable for *in vivo* animal studies due to increased depth penetration, it is especially challenging to find appropriate FRET pairs and other fluorophores with enough spectral separation. In addition, NIR fluorophores often possess prohibitively short lifetime, which presents extra challenge for the lifetime multiplexing approach.

Recent development of dark quenchers,^33–35^ which do not produce inherent fluorescence emission but still undergo quenching via several mechanisms, including FRET and static quenching, has been harnessed to mitigate high signal crosstalk and allow multiplexing FRET pairs with other fluorophores. These quenchers have been primarily designed for visible fluorophores, but some quenchers such as black hole quencher-3 (BHQ-3) and SiNQs^34^ have been developed for longer-wavelength probes. However, the application of these quenchers is limited due to their relatively narrow spectrum of absorbance. Conversely, IRDye QC-1 (QC-1; Li-Cor, Inc.) is a recently developed dark quencher with a much broader absorbance spectrum that encompasses the NIR and visible range.^36, 37^ To date, QC-1 has been used as a quencher in activatable probes in imaging of antibody internalization,^38, 39^ enzymatic activity,^40^ and as a marker for photoacoustic imaging.^41, 42^ To the best of our knowledge, this is the first report of QC-1 complete characterization as an acceptor for NIR FLI-FRET *in vivo* imaging in live intact animals. First, we established the spectral characteristics of QC-1 using hyperspectral single-pixel and wide-field MFLI *in vitro* imaging. Second, the uptake of Tf-QC-1 conjugates into breast cancer cells was analyzed at multiscale using confocal microscopy, *in vitro* NIR FLIM FRET microscopy and, finally, *in vivo* NIR MFLI-FRET imaging. Last, we report on the multiplexed imaging of NIR-labeled Tf FRET pair (Tf-AF700/Tf-QC-1) uptake in tumors concurrently with the metabolic marker 2-DG. Overall, this body of work ascertains the utility of QC-1 as an effective FRET acceptor for the monitoring of receptor-ligand interactions, *i.e.* drug-target engagement, *in vitro* and *in vivo* with the benefit of opening a wide range of opportunities to monitor multiplexed ligands or antibodies biodistribution and internalization using MFLI-FRET whole body imaging in preclinical studies. Importantly, multiplexing of NIR FRET with 2-DG imaging provides invaluable insight into the drug binding and metabolic drug response in real time to optimize tumor drug delivery.

## Results and Discussion

### Monitoring target engagement and metabolic levels in tumor xenografts in live mice

The ability to measure the binding and internalization of targeted therapeutics into tumor cells in combination with the detection of metabolic levels in an undisturbed tumor environment could provide fundamental information regarding molecular profile, delivery efficacy and drug response of tumor xenografts. In this study, we capitalize on TfR-Tf binding to quantify target engagement and internalization of ligand-receptor complexes. Tf, a major blood serum glycoprotein, is responsible to transport iron to every cell in the body. Therefore, Tf has been used as a carrier for targeted drug delivery due to the upregulation of its respective receptor, *i.e.* TfR, in pathological tissues, including cancer. ^43, 44^ In a first attempt of simultaneous monitoring metabolic imaging and TfR-Tf internalization in tumor, we performed MFLI imaging to measure Tf-TfR engagement using Tf-AF700/Tf-AF750 FRET pair and 2-DG fluorescence signal, a NIR-labeled metabolic marker, in a mouse bearing T47D breast cancer xenograft. Since AF750 excitation and emission spectra overlap considerably with those of IRDye 800CW (Suppl. Fig. 1A-B), MFLI imaging was performed sequentially as shown in Fig. 1A. Firstly, the animal was tail-vein injected with 2-DG and imaged 24 h post-injection. Only after that, the NIR-labeled Tf FRET pair at A:D = 2:1 was tail-vein injected and imaged 1 h post-injection. Although 2-DG could be detected in many organs, it accumulated predominantly in the tumor (Fig. 1B) because of higher level of glycolysis compared to normal tissues. Tumor and liver displayed elevated levels of FD%, indicating binding and internalization of NIR-Tf into TfR-enriched tissues (Fig. 1C). In contrast, as expected, the urinary bladder showed very low levels of FD%, suggesting that fluorescence signal is due to excreted Tf-AF700 degradation products.^22–24^ Although successful, this study was performed using a sequential imaging protocol that increases workload, can lead to some registration errors between the spatial mapping of the biomarkers, and may introduce biological bias due to the dynamic nature of tumor biology. Therefore, we investigated the potential of using dark quencher QC-1 as an alternative FRET acceptor to enable multiplexed imaging in the same animal and concurrent imaging session (Suppl. Fig. 1C).

**Figure 1.**
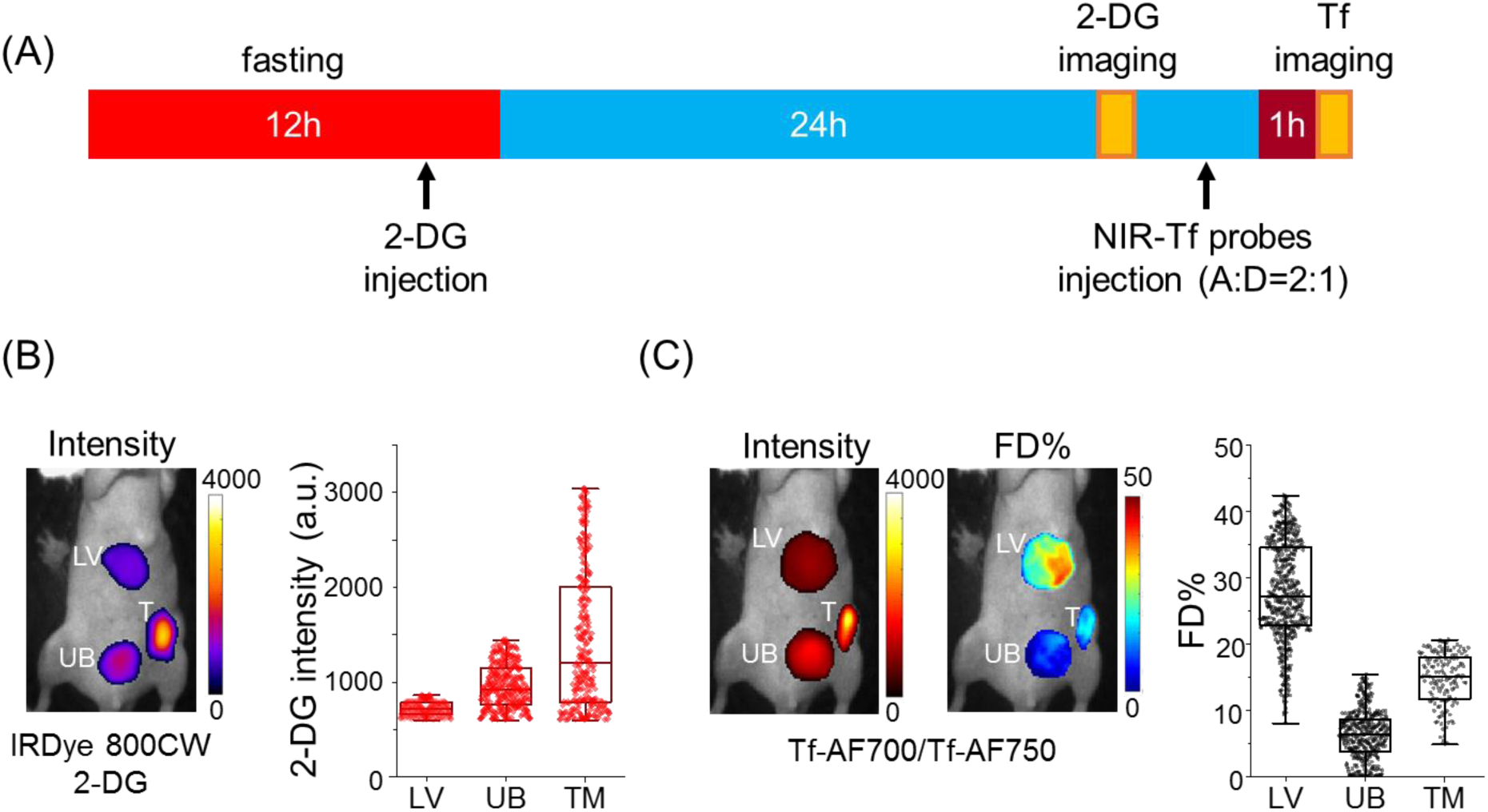
Sequential multiplexed MFLI FRET and metabolic in vivo imaging: (A) Imaging protocol includes fasting prior to IRDye 800CW 2-DG (2-DG) injection and imaging, followed by NIR-FRET pair injection and imaging. (B) Representative image of 2-DG fluorescence intensity (left panel) and respective graph displaying distribution of 2-DG signal per ROI pixel (right panel) of mouse liver (LV), tumor xenograft (TM) and urinary bladder (UB). (C) Left panel, Tf-AF700 intensity (total Tf, including bound and unbound) and FD% (bound Tf) map of liver, tumor and urinary bladder in mouse sequentially injected with 2-DG and Tf-AF700/Tf-AF750 and imaged with MFLI at 24 h and 1 h post-injection, respectively. Right panel shows distribution of FD% signal per ROI pixel. Rectangle box indicates 25%-75% pixel values, horizontal line indicates median value and vertical line indicates range within 1.5 quartile.

### Characterization of IRDye QC-1 as a NIR dark quencher FRET acceptor using hyperspectral single-pixel imaging

We investigated the spectral characteristics of the FRET pair Alexa Fluor 700 (AF700)/QC-1 using a hyperspectral single-pixel imager. To this end, we performed an antibody binding FRET multi-well assay, in which samples are composed of AF700 conjugated to murine IgG primary antibody (IgG-AF750) and goat anti-mouse secondary antibody conjugated to QC-1 (anti-IgG-QC-1) or AF750 (anti-IgG-AF750). While keeping the donor concentration constant at 50 µg/mL, the samples contained or not, increasing concentration of respective acceptor at various A:D ratios (0:1, 1:1, 2:1 and 3:1). A hyperspectral single-pixel imaging system with lifetime capabilities was used to visualize donor and acceptor emission spectra as well as quantify FRET lifetime parameters. Technically, upon excitation at 695 nm (0.5 s per illumination pattern; ∼5 min acquisition for multi-well plate), fluorescence emission was detected via a spectrophotometer and distributed into a 16-channel PMT detector using a wide-field lifetime-based hyperspectral single-pixel imaging platform. The acquired emissions were reconstructed to time-domain datasets to further least-square-fit each pixel for lifetime quantification. ^45^

Figure 2 displays the continuous wave (CW) intensity reconstructions (Fig. 2A, D) and intensity quantifications (Fig. 2B-C & E-F) for each A:D ratio and NIR FRET pair in multi-well plate samples. As expected, upon excitation at 695nm, the wells containing PBS (Fig. 2A, D: b1, c1) and QC-1 conjugate (Fig. 2D: a3, b3, c3) showed no emitted fluorescence signal. In contrast, wells containing AF700 and AF750 or QC-1 at A:D ratios of 0:1 (a1), 1:1, 2:1 and 3:1 (a2, b2, c2) displayed a decreasing AF700 fluorescence intensity as the AF750 (Fig. 2A-B) or QC-1 (Fig. 2D-E) concentrations increased. Importantly, acceptor bleed-through was observed for AF750 (Fig. 2B) but not for QC-1 (Fig. 2E). The maximum fluorescence intensity of AF700 signal was detected between 720 and 738 nm (Fig. 2B, E) as opposed to the maximum intensity between 747 and 765 nm for AF750 only wells (Fig. 2B).

**Figure 2.**
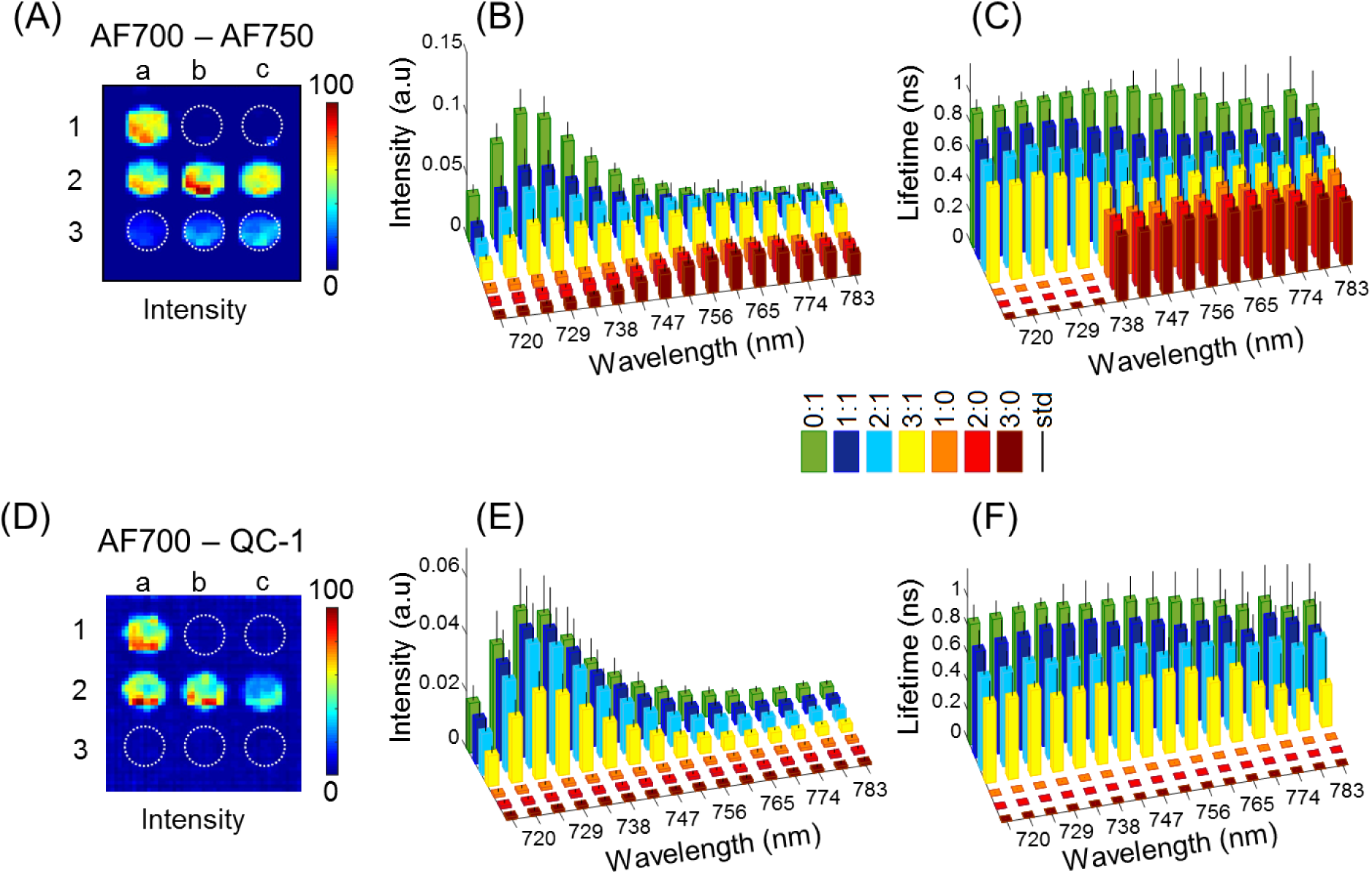
Hyperspectral characterization of QC-1 as a FRET acceptor. (A) Reconstructed spectrally resolved CW intensity spatial map of AF700 donor and AF750 acceptor at the 725 nm wavelength. Spectrally resolved mean intensity (B) and lifetime (C) distributions for each well per wavelength and A:D ratio for AF700/AF750 FRET donor/acceptor sample. (D) Reconstructed spectrally resolved CW intensity spatial map of AF700 donor and QC-1 acceptor at the 725 nm wavelength. (E) Mean intensity and (F) lifetime per wavelength and A:D ratio for AF700/QC-1 FRET donor/acceptor sample. For both (A) and (D), the layout of the A:D ratios on the plate is as follows: a1-0:1, b, c1-0:0, a2-1:1, b2-2:1, c2-3:1, a3-1:0, b3-2:0 and c3-3:0. Error bars represent standard deviation.

The reconstructed fluorescence lifetime maps show that the lifetime was reduced in all FRETing samples (Fig 2C, F). Decreasing lifetime correlated with increasing A:D ratios 1:1, 2:1 and 3:1, due to elevated FRET-mediated donor quenching events as the acceptor concentration increases ^14^. The lifetime value for AF700 donor alone (A:D ratio = 0:1) remains constant across wavelengths (∼0.8-0.9 ns), as lifetime is independent of fluorescence intensity. Moreover, AF700 lifetime displays higher standard deviations at the higher wavelengths, when donor signal is reduced (Fig. 2C, F). The AF700/AF750 FRET pair displayed decreased lifetime of each FRETing sample with increasing wavelengths (Fig. 2C). Contribution from the acceptor (AF750) fluorescence, which exhibited a shorter lifetime (∼0.5 ns), leads to significant reduction in lifetime values of FRETing AF700/AF750 samples and could be confirmed by the presence of AF750 fluorescence in late spectral channels (> 738 nm). In contrast, the lifetime of each A:D ratio of FRETing AF700/QC-1 sample remained constant for all spectral channels, even though the standard deviation was greater at longer wavelengths due to reduced fluorescence intensity that could affect reliability of the calculation. QC-1’s quencher properties minimized the spectral crosstalk, and thus improved the accuracy of the lifetime result, allowing for direct quantification of AF700 FRETing donor fraction and mean lifetimes even in the absence of specific emission filters or spectral unmixing. In summary, the hyperspectral imaging analysis demonstrated the superiority of QC-1 as a FRET acceptor when compared to AF750.

### MFLI-FRET analysis of AF700-QC-1 FRET pair

Next step was testing QC-1 as the acceptor in an antibody binding FRET assay using wide-field MFLI imaging platform, which includes emission filters to selectively collect donor fluorescence without being contaminated by the excitation light or AF750 crosstalk, as described in Experimental Methods. Similar to the hyperspectral experiment (Fig. 2), we used murine AF700-IgG as the donor and AF750 or QC-1 conjugated to goat anti-mouse secondary IgG as the acceptor in FRET reaction. The FRET pairs were mixed in A:D ratios of 0:1, 0.5:1, 1:1, 2:1, 3:1 with the donor concentration constant at 50µg/mL (Fig. 3A-B). Although fluorescence decays at A:D=2:1 look identical (Fig. 3C), FD% quantification plotted against the A:D ratio (Fig. 3D) shows that dark acceptor QC-1 significantly outperforms the emitting acceptor AF750 (Spearman correlation coefficient p=0.0167). Thus, we confirmed QC-1 as a superior FRET acceptor when using MFLI wide-field imaging approach.

**Figure 3.**
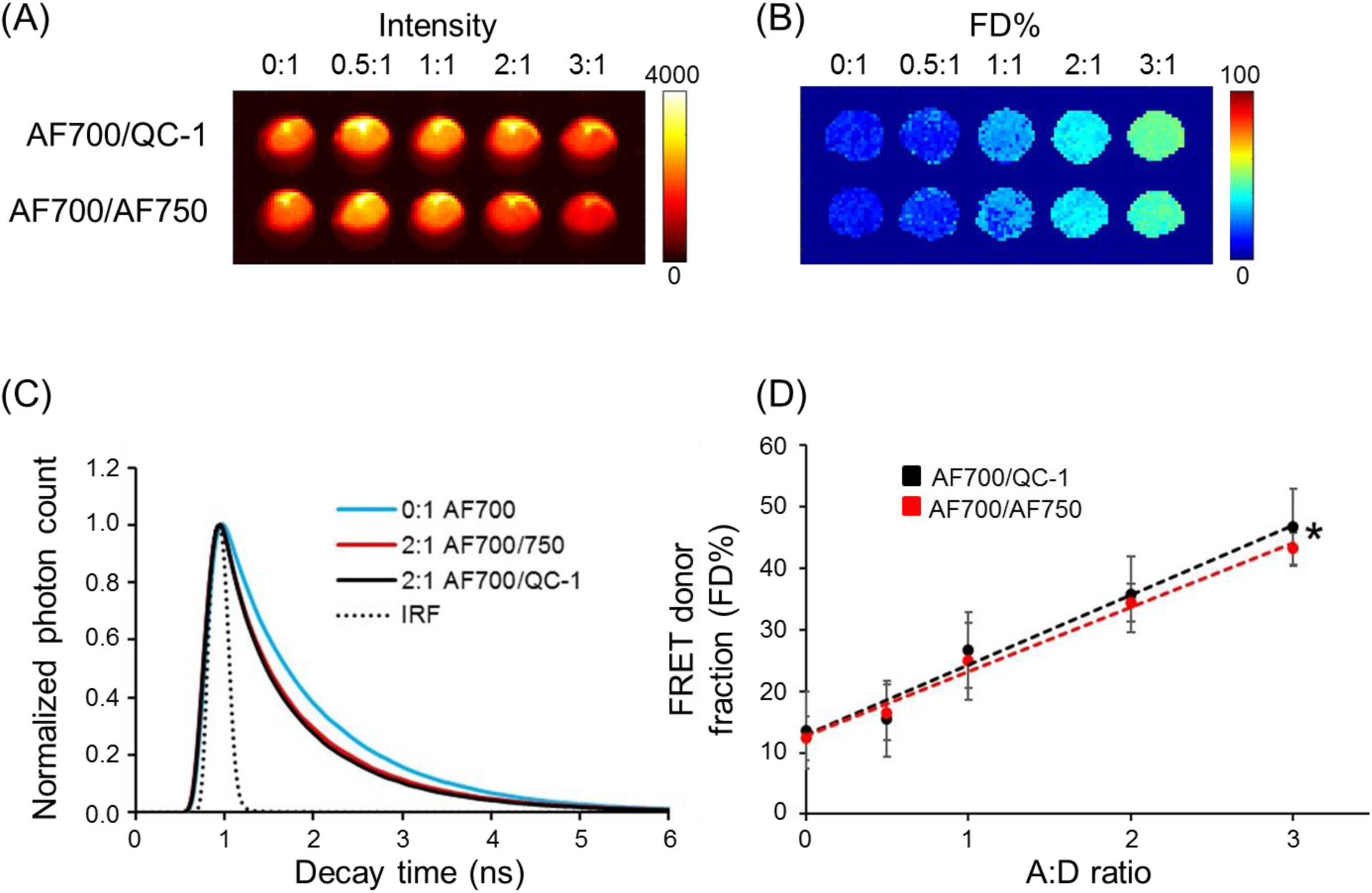
Comparison of AF700/AF750 vs. AF700/QC1 FRET pairs using wide-field MFLI. Fluorescence intensity (A) and FD% (B) maps of AF700/AF750 and AF700/QC1 FRET samples. (C) Donor fluorescence lifetime decay of AF700/AF750 (red line) and AF700/QC1 (black line) samples with A:D ratio 0:1 and 2:1; IRF is instrument response function. (D) Quantification of FD% vs A:D ratio for AF700/AF750 (red) and AF700/QC (black) antibody binding FRET assay. Data presented as a mean ± standard deviation. Spearman correlation between two linear regression lines p-value (two-tailed) is 0.0167 (asterisk). Statistical analysis is described in Supplementary Table 1.

### Tf-AF700 and Tf-QC-1 internalize into breast cancer cells in a similar manner

To verify that Tf-QC-1 undergoes receptor-mediated endocytosis similarly to previously characterized Tf-fluorophore conjugates in relevant biological settings, we subjected T47D human breast cancer cells to Tf uptake imaging assay by incubation with 40µg/mL Tf-AF700 or Tf-QC-1. This breast cancer cell line overexpresses TfR and estrogen receptor and represents the most common type of breast cancer - luminal A.^46^ The degree of colocalization of Tf-AF700 vs. Tf-QC-1 with TfR, which is a major early and recycling endocytic marker,^47^ was assessed in Tf-treated cells and then processed for immunostaining using antibodies against human Tf and TfR (Fig. 4A). In case Tf-QC-1 binds TfR less efficiently, the degree of TfR/Tf-QC-1 colocalization would be less in comparison to that of TfR/Tf-AF700. Z-stacks were collected on confocal microscope and analyzed using Imaris software to calculate Pearson’s correlation coefficient (Fig. 4A). Fig. 4B shows that there is no significant difference (Suppl. Table 2, p=0.089) between TfR and Tf-AF700 or Tf-QC-1colocalization. These results confirmed that both ligands are internalized and trafficked in a similar manner via receptor-mediated endocytic pathway.

**Figure 4.**
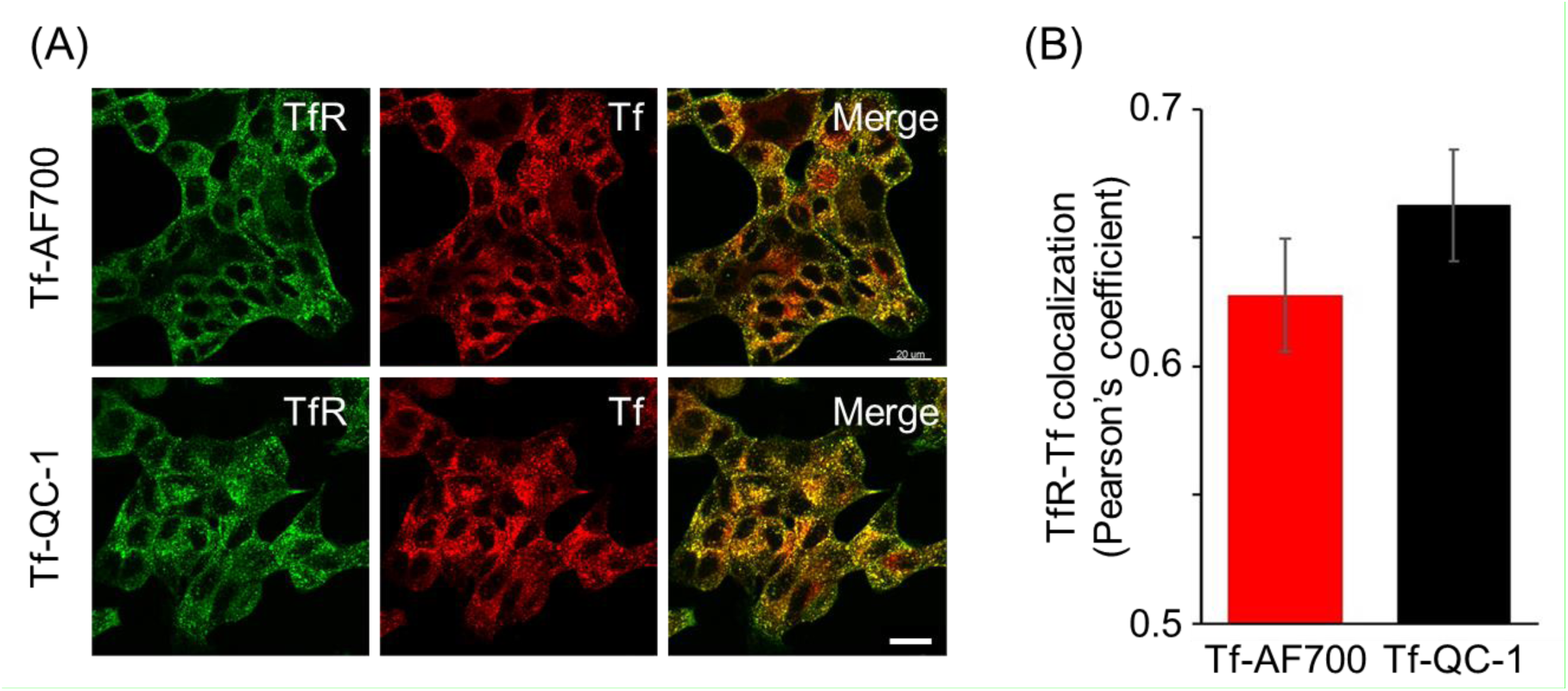
Comparison of Tf-AF750 and Tf-QC-1 internalization and trafficking in cancer cells. (A) Maximum intensity projections of z-stacks consisting of 10-12 optical slices of T47D human cancer cells loaded with Tf-AF700 or Tf-QC-1 and processed for anti-TfR (green) and anti-Tf (red) immunostaining. Scale bar = 20 µm. (B) Quantification of TfR and Tf colocalization analysis using Pearson’s correlation coefficient throughout the entire z-stack using Imaris imaging analysis software. Data presented as a mean (n=5), error bars represent standard deviation, p=0.089 (Suppl. Table 2).

### QC-1 as an acceptor in FRET assays using NIR FLIM and MFLI imaging

In order to validate QC-1 as an acceptor for FRET using FLI FRET microscopy (FLIM FRET), we performed Tf uptake assays in T47D cells using Tf-AF700/Tf-AF750 and Tf-AF700/Tf-QC-1 ligands at various A:D ratio. After washing off the ligands and fixing the cells, the samples were imaged using NIR FLIM system and analyzed with SPCImage (Becker & Hickl). NIR FLIM was performed in Zeiss LSM880 microscope using the descanned detection pathway via direct connection and a Titanium: Sapphire laser (Chameleon) as a single-photon excitation source (Suppl. Fig. 2). An excitation wavelength of 690nm was selected for cells incubated with Tf-AF700 (donor specimen), Tf-AF700 and Tf-AF750 (donor plus acceptor sample) or Tf-AF700 and Tf-QC-1 (donor plus dark quencher acceptor); at this wavelength, cells incubated with only Tf-AF750 showed negligible intensity levels. The amount of donor photobleaching was monitored by measuring intensity changes over time. Donor photobleaching levels were reduced (<5%) at the 690nm excitation power level used in NIR FLIM-FRET experiments

Fig. 5A shows that both FRET pair ligands demonstrate the same pattern of distribution within the cells. Likewise, both samples had significantly shorter lifetime compared to single donor sample in fluorescence decay graphs (Fig. 5B) and frequency distribution (Fig. 5C), as well as linear trend of FD% with increasing A:D ratio (Fig. 5D). However, in contrast to antibody binding experiment (Fig. 3), a small but statistically significant decrease in FD% levels was observed in Tf-AF700/Tf-QC-1 cell samples when compared to that in Tf-AF700/Tf-AF750 cells.

**Figure 5.**
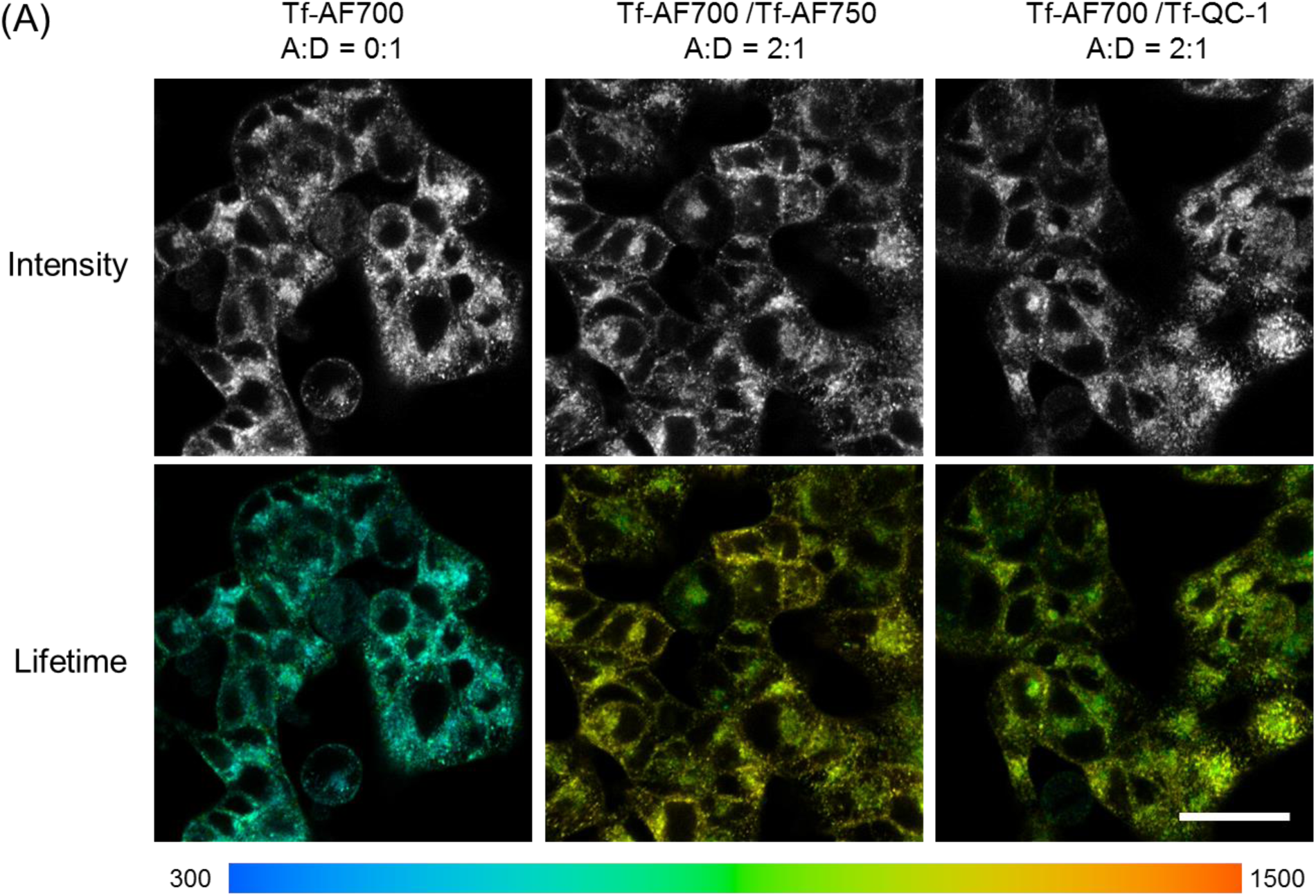
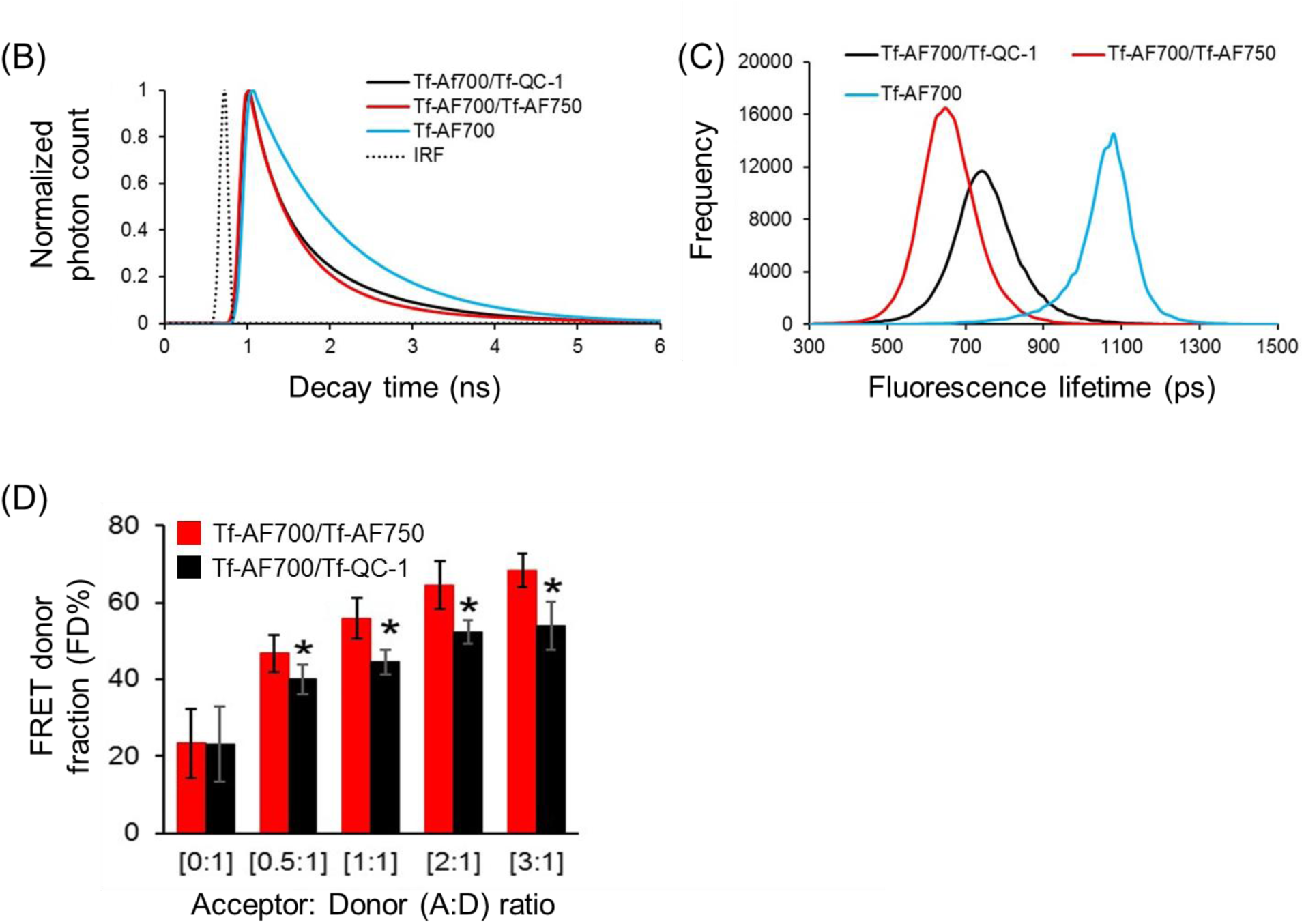
NIR FLIM FRET assay in cancer cells. (A) NIR FLIM TCSPC data. The representative images of fluorescence intensity and lifetime map (τ_*m*_) in T47D cells treated with Tf-AF700 (A:D=0:1), Tf-AF700 and Tf-AF750 (A:D=2:1) or Tf-AF700 and Tf-QC-1 (A:D=2:1); pseudo-color range= 300-1,500 ps. Both fluorescence intensity and lifetime distributions show punctate endocytic structures containing TfR-Tf complexes. Scale bar= 50 µm. (B) Representative fitting curves and IRF, the fluorescent lifetime decay in the single and double-labeled cells was determined by comparing the fitting of the decay data using both single- and double-exponential decay models. (C) Comparison of fluorescent lifetime distribution in T47D cells treated with Tf-AF700 (A:D=0:1), Tf-AF700 and Tf-AF750 (A:D=2:1) or Tf-AF700 and Tf-QC1 (A:D=2:1). (D) Comparison of FRET levels in T47D cells treated with Tf-AF700/Tf-AF750 or Tf-AF700/Tf-QC1 FRET pairs at various A:D ratios. N=10, error bars represent standard deviation, p < 0.05 (Suppl. Table 3).

The first *in vivo* validation of QC-1 as a FRET acceptor was performed in live anesthetized mice implanted with Matrigel plugs containing cancer cells preloaded with Tf-AF700/Tf-AF750 or Tf-AF700/QC-1 at A:D ratio 0:1, 1:1 and 2:1 (Suppl. Fig. 3A). The goal of this experiment was to compare the relationship between FD% vs A:D ratio using these two NIR FRET pairs as they are imaged through living tissues. With such experimental design, the donor signal was localized strictly to the plugs. However, because of the different size of injections and the localization of the plug on the animal body, significant variation in fluorescence intensity was detected across the different A:D ratio and FRET pairs in Matrigel plugs. Nevertheless, due to lifetime’s independence from fluorescence intensity, FRET levels in both mice were very comparable, showing linear trend with increasing A:D ratio. Again, less FD% in Tf-AF700/Tf-QC-1 plugs were observed (Suppl. Fig. 3B). In summary, these results suggest that Tf-QC-1 acts as an suitable FRET acceptor in both NIR FLIM microscopy (Fig. 4) and macroscopy (Suppl. Fig. 3).

From the previous experiments, we determined that Tf-QC-1 can bind TfR at similar levels to Tf-AF conjugates and TfR-Tf-QC-1 receptor-ligand complexes can be transported adequately via the endosomal pathway in T47D cells. Based on the antibody binding assays, QC-1 NIR FRET performance was just as good as “classical” acceptor AF750 with the advantage of reduced bleed-through levels. However, Tf-AF700/QC-1 pair consistently showed relatively less NIR FRET compared to Tf-AF700/Tf-AF750 in 2D cell culture using FLIM FRET as well as in living intact animals using wide-field MFLI imaging. The most likely reason for that is the difference in Förster distance (R0) between AF700-AF750 (R0= 78.1Å) and AF700-QC-1 (R0= 69Å) FRET pairs (Suppl. Fig. 4). Reduced R0 decreases overall FRET efficiency and putative impact of “crowding” effect.^48^ One should have in mind that *in vitro* antibody binding assay is done in an ideal homogeneous solution, which is far from real life scenario of cells with different ligand local concentration gradient and receptor density. The reduced Förster distance of AF700/QC-1 may decrease the possibility of FRET between two neighboring Tf-TfR complexes, which could contribute to reduced FD% levels in cells loaded with Tf-AF700/Tf-QC-1 FRET pair.

### MFLI-FRET in vivo imaging of liver and urinary bladder

The ultimate QC-1 validation was performed *in vivo* by imaging several mice intravenously injected with Tf-AF700/Tf-AF750 or Tf-AF700/Tf-QC-1 probes at A:D ratio 2:1. MFLI imaging was performed at 2 h, 6 h and 24 h post-injection to monitor NIR-labeled Tf biodistribution and target engagement. We focused on the liver, a main organ for pharmacokinetic studies, which is also a main site for iron metabolism with highly elevated TfR levels. Urinary bladder, an important excretion organ, was used as a negative control since it is expected to accumulate degradation products and free dye. Substantial amount of FRET, representing the fraction of internalized NIR-Tf from injected probe, was detected in the liver, but not in the bladder, as shown in representative images of mice displaying Tf-AF700 donor fluorescence intensity and FD% quantification (Fig. 6A-B). FD% levels in the liver remained at the same low level in the control animal injected with donor only during time course, whereas FD% in the livers of mice injected with either FRET pair increased up to three-fold above the control level (Fig. 6C). However, the dynamics of Tf internalization in the liver represented as FD% differed in Tf-AF750 and Tf-QC-1 groups (Fig. 6C). Whereas FD% in the Tf-AF700/Tf-AF750 injected mice peaked at 6 h p.i., the Tf-AF700/Tf-QC-1 group demonstrated a trend to remain at the flat level during the 24 h course of imaging. Overall, QC-1 proved to be an efficient acceptor for FRET pair both for *in vitro* and *in vivo* applications using lifetime-based imaging approaches.

**Figure 6.**
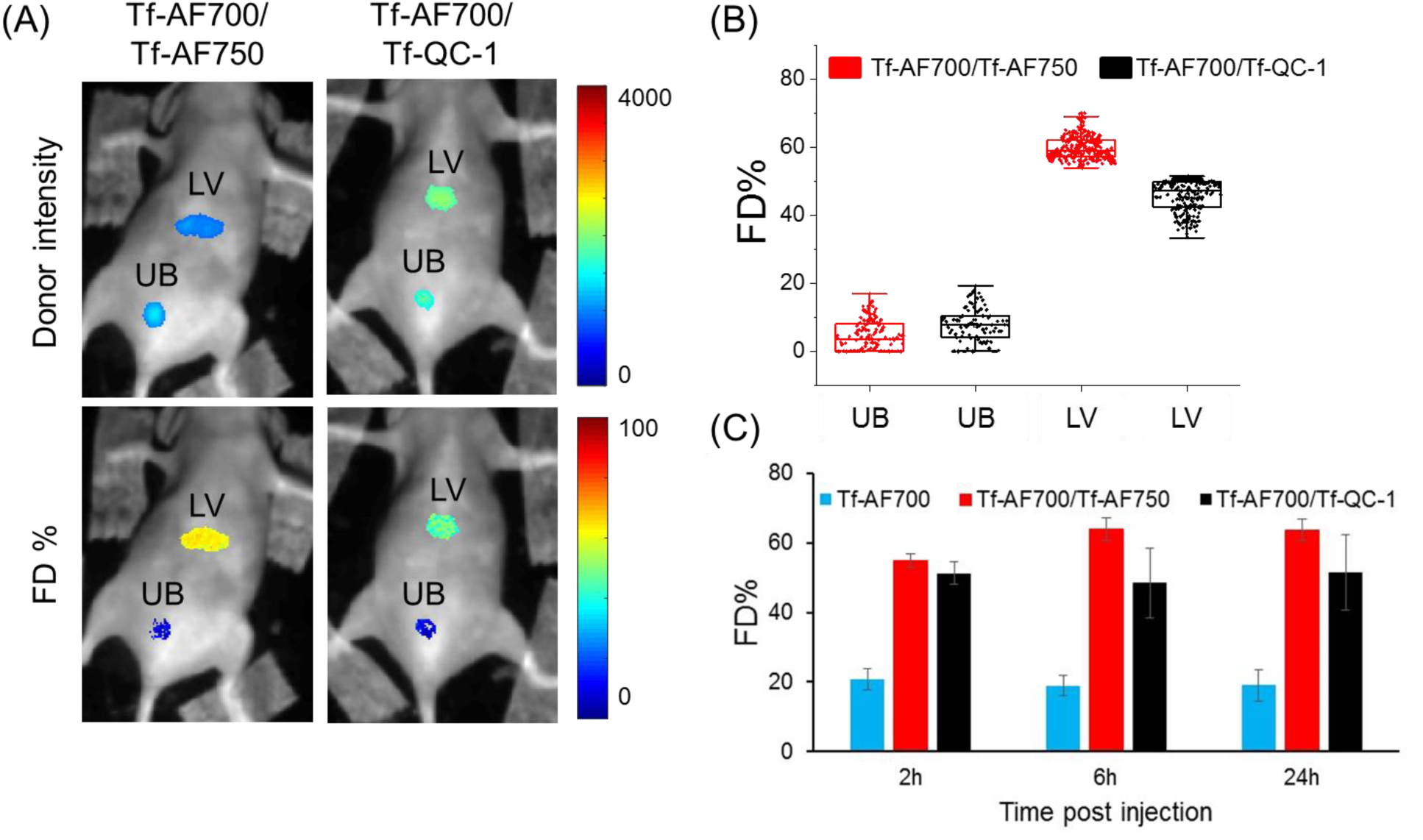
Whole-body MFLI FRET *in vivo* imaging. (A) Representative images of donor intensity (total Tf, including bound and unbound) and FD% (bound Tf) map of liver (LV) and urinary bladder (UB) in mice tail-vein injected with 40µg/mL Tf-AF700 and 80µg/mL Tf-AF750 or Tf-QC-1 (A:D= 2:1) and imaged using MFLI imager at 24 h p.i. (B) Graph displaying distribution of FD% signal per ROI pixel in livers and bladders of shown mice at 24 h p.i., as described in Fig. 1B-C legend. (C) Time course of Tf internalized in livers of 4 mice injected with Tf-AF700/Tf-AF750 and 3 mice injected with Tf-AF700/Tf-QC-1 from two independent experiments. Error bars represent standard deviation.

### Multiplexed MFLI-FRET in vivo whole-body imaging

Finally, we tested QC-1 for multiplexing by simultaneous imaging Tf-AF700/Tf-QC-1 and 2-DG, a NIR-labeled glucose metabolic marker, in mice bearing T47D tumor xenografts (Fig. 7). Mouse 1 (M1) had a large aggressively growing tumor of 455.5 mm^3^ volume, whereas mouse 2 (M2) tumor had a much smaller size of 98.6 mm^3^. The ability to simultaneously image a NIR FRET pair together with NIR 2-DG probe was possible only because of QC-1 as acceptor, since AF750 excitation and emission significantly overlap with those of IRDye 800CW (Suppl. Fig. 1C). Here, we successfully visualized tumors and urinary bladder (Fig. 7B) as well as liver (Suppl. Fig. 3A) at 24 h p.i. by sensing the accumulated 2-DG probe and detectable FRET signal from internalized Tf-AF700/Tf-QC-1. Fig. 7A displays the imaging protocol used for this experiment, in which animals were subjected to fasting for improved visualization of 2-DG signal, followed by two staggered injections of 2-DG and NIR Tf FRET pair and a single imaging session using wide-field MFLI. Fig. 7B shows that both tumors are easily detected by 2-DG, which specifically accumulates in metabolically active tissues with elevated level of glycolysis and glucose transporter 1 (GLUT1).^49^ However, M1 mouse displayed significantly higher level of 2-DG intensity in the tumor than M2 mouse (Fig. 7C) indicating that the large aggressively growing tumor exhibited much higher rate of glycolysis. In contrast, there was inverse correlation between 2-DG fluorescence intensity, Tf fluorescence intensity (total free and internalized ligand) in the tumors, and FD% (fraction of receptor bound and internalized ligand) (Fig. 7C-D and 7F). Unlike M1 tumor, smaller M2 tumor accumulated much less 2-DG and total Tf but exhibited significant amount of Tf-TfR engagement represented as FRET signal. Stacked bar charts of 2-DG and Tf pixels frequency of both tumors show this inverse relationship in a compelling manner (Fig. 7E and 7G). To eliminate the potential variability of amount of injections, we normalized both 2-DG and FRET values to the corresponding values in the liver. Nevertheless, the difference in 2-DG fluorescence intensity between the tumors was still dramatic (Suppl. Fig. 3). Importantly, histological analysis confirmed that larger tumor M1 had almost background level of Tf staining whereas M2 tumor displayed significant number of Tf-positive cells (Fig. 8). Both tumors however exhibited similarly high levels of TfR across the surface (Fig. 8c and 8h) which is consistent with our previous observation. ^22^

**Figure 7.**
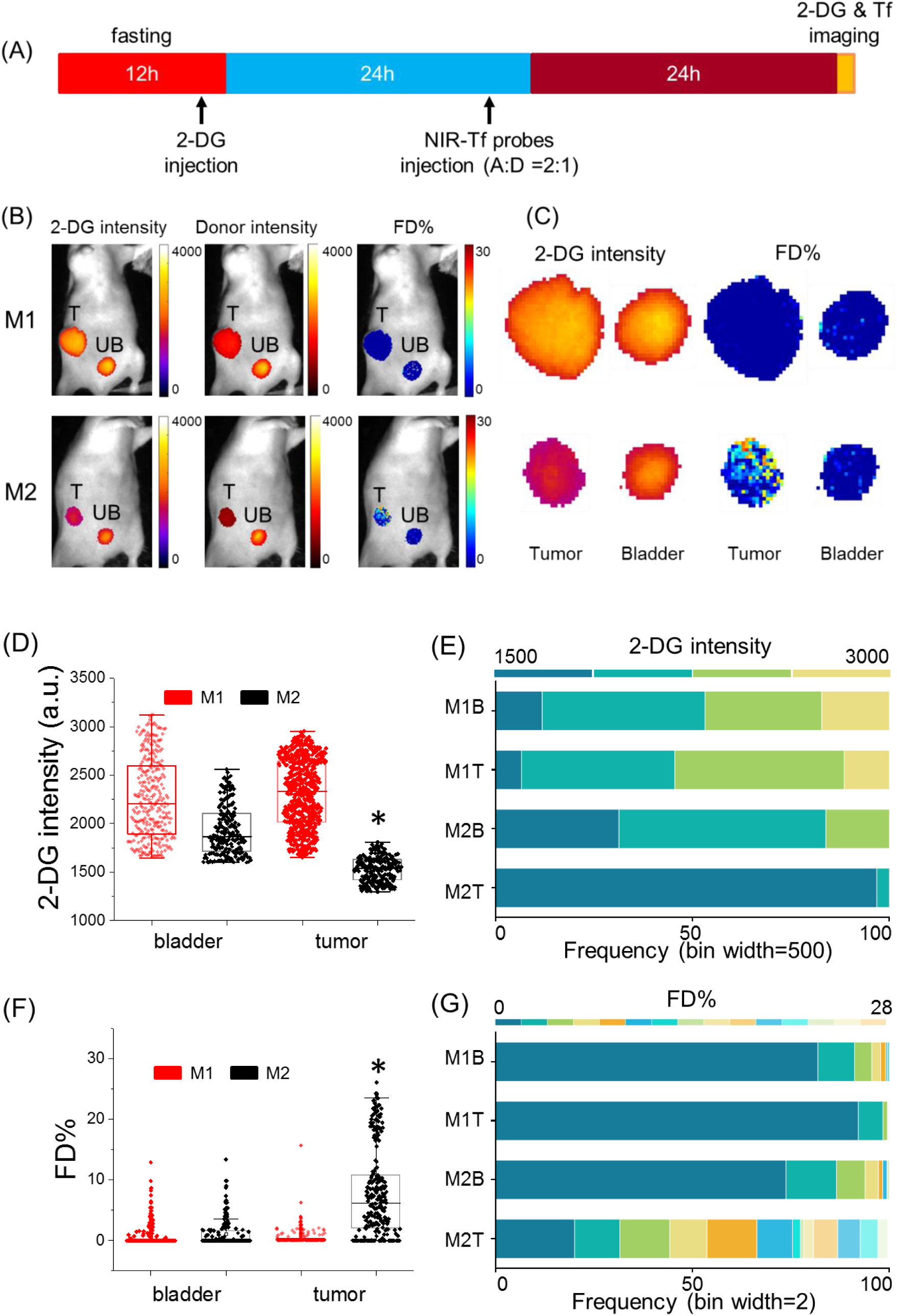
Multiplexed MFLI FRET in vivo imaging: (A) Imaging protocol including fasting and 2-DG and NIR-FRET pair injections. (B) Representative images of 2-DG and Tf-AF700 intensity and FD% map of tumor and urinary bladder in mice simultaneously injected with 2-DG and Tf-AF700/Tf-QC-1 and imaged with MFLI FRET at 24 h post-Tf injection. 2-DG mask was used to determine Tf ROI with the same number of pixels. (C) Magnified pixels of 2-DG intensity and FD% in tumors. (D) Distribution of 2-DG fluorescence intensity per ROI pixel in tumors and bladders. (E) Stacked frequency bar chart of 2-DG fluorescence intensity, bin width = 500 a.u. (F) Distribution of FD% signal per ROI pixel in tumors and bladders. (G) Stacked frequency bar chart of Tf FD%, bin width = 2. Asterisks indicate p < 0.05 (Suppl. Table 4).

**Figure 8.**
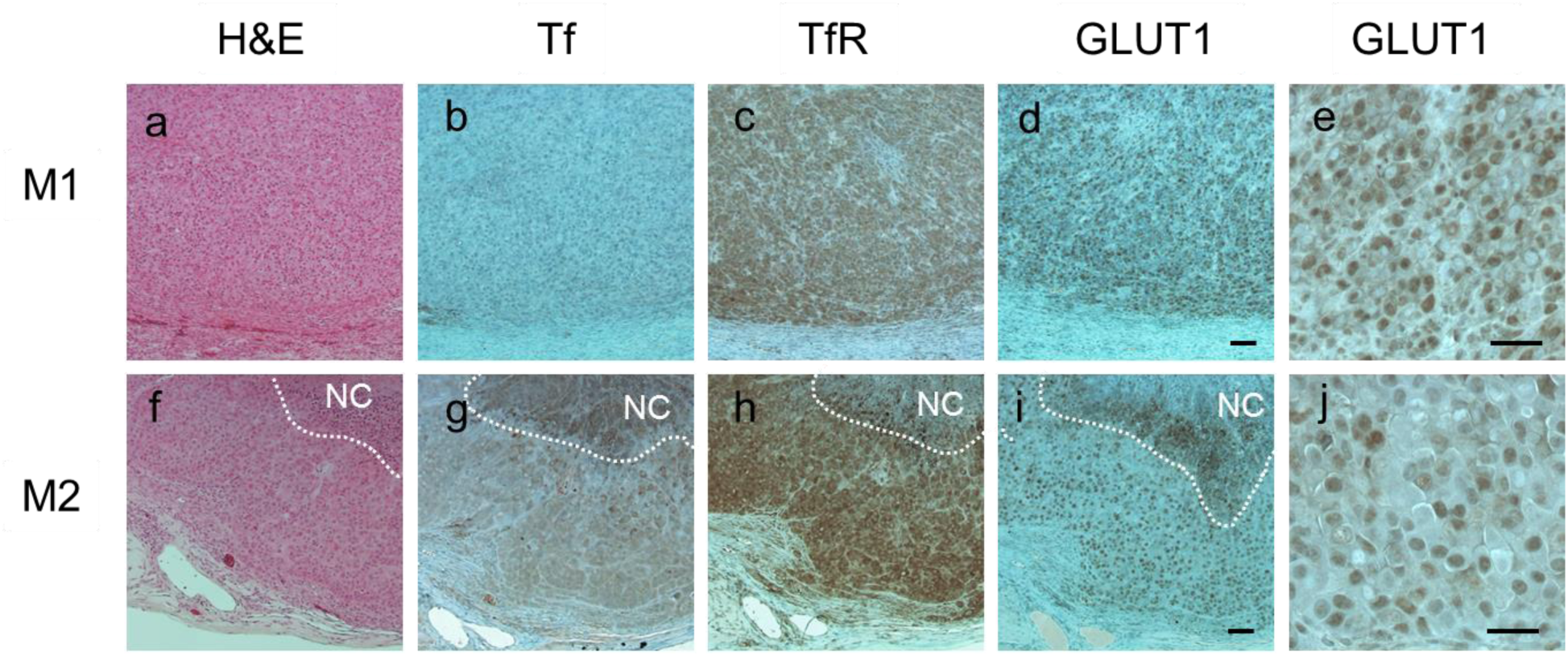
Histological analysis of consecutive tumor sections from M1 (a-e) and M2 (f-j) T47D xenografts processed for hematoxylin and eosin staining (a & f), anti-Tf (b & g), anti-TfR (c & h) and anti-GLUT1 (d & i, e & j (40x) immunohistochemical staining. NovaRED was used as peroxidase substrate (brown color), sections were counterstained with methyl green. White dashed lines delineate necrotic core (NC). Scale bar= 100 µm.

Of importance, immunohistochemistry (IHC) staining for glucose transporter 1 (GLUT1) matched 2-DG imaging data: M1 tumor had visibly higher staining density of GLUT1 and, unlike M2 tumor with predominantly cytoplasmic GLUT1 staining (Fig. 8 i, j), displayed actual membranous localization (Fig. 8d, e). It has been reported that GLUT1 tends to localize to the cytoplasm in human breast cancers.^50^ What makes it especially relevant is that GLUT1 expression is upregulated by hypoxia-induced factor 1 (HIF-1) and is highly correlated with poor prognosis and survival in most solid tumors.^51, 52^

In summary, these results demonstrate for the first time the utility of QC-1 for multiplexed NIR FRET-MFLI imaging in living intact mice carrying tumor xenografts. 2-DG is useful for delineation of tumor margins thus eliminating the reliance on donor intensity-based ROI selection. Unexpectedly, we observed 2-DG potential to predict the tumor resistance to efficient intracellular ligand delivery as indicated by Tf FD% vs 2-DG relationship. It is probable that aggressively growing tumors characterized by poor blood perfusion, stiff extracellular matrix and thus near impenetrable for drug delivery,^53, 54^ also possess highest levels of glycolysis. Our experience with MFLI FRET imaging of Tf-TfR target engagement in such tumors represented by triple negative breast cancer MDA-MB 231 xenografts confirms this trend: FD% comparable to the “normal” T47D xenografts was detected only at 48 h p.i. (Suppl. Fig. 6A-C). IHC analysis also demonstrated higher level of expression of glucose transporter GLUT-1 in MDA-MB 231 vs T47D tumors (Suppl. Fig. 6D). These results are consistent with the analysis of metabolic markers in basal type vs non-basal type human breast cancer samples.^55^

Multiplexing *in vivo* imaging of Tf-AF700/Tf-QC-1 FRET signal with that of 2-DG demonstrates a vast potential for simultaneous monitoring of metabolic response and intracellular drug delivery using MFLI-FRET. In fact, the reduction of metabolic activity in non-small cell lung cancer by ^18^F-FDG PET imaging has been reported in clinical study as an early prediction of therapeutic response to nivolumab.^56^ Our technology offers similar but much straightforward and accessible method with the additional benefit of multiplexing. With the advancement of new NIR fluorophores and probes on the market, it is a matter of time to develop a protocol for robust monitoring of polytherapy drug delivery/metabolic response in preclinical studies. That is when dark quencher QC-1 as an acceptor becomes practical and, in many cases, the only option for efficient multiplexed FRET sensing. This multiplexed approach is poised to improve significantly the ability to perform drug delivery optimization in pre-clinical drug development research.

## Conclusion

In this study, we fully characterized dark quencher QC-1 as NIR FRET acceptor using hyperspectral, NIR FLIM and MFLI imaging platforms, *in vitro* and *in vivo*. We demonstrated that QC-1 offers a possibility to become an essential tool for simultaneous quantification of target engagement and monitoring tumor metabolic status. This multiplexed approach is crucial for a non-invasive drug delivery assessment and metabolic response, as it provides a robust means to drastically improve the efficacy of drug development in preclinical studies.

## Experimental Methods

### Ligand labeling

Human holo Tf (Sigma) was conjugated to AF700 or AF750 (Life Technologies) through monoreactive N-hydroxysuccinimide ester to lysine residues in the presence of 100 mM Na bicarbonate, pH 8.3, according to manufacturer’s instructions.^14^ The probes were purified by desalting columns and dialysis. The degree of labeling of the probes was assessed by spectrophotometer DU 640 (Beckman Coulter, Fullerton, CA, USA). The average degree of labeling was no more than 2 fluorophores per Tf molecule. Custom Tf-QC-1 conjugation was performed by Li-Cor (Lincoln, NB, USA) with average dye to protein ratio of 3. All ligands were normalized to concentration 1 mg/mL in phosphate-buffered saline pH 7.6 and filter sterilized.

### Hyperspectral imaging

AF700 (AF700; MG129, Thermo Fisher Scientific, MA) conjugated to murine IgG primary antibody was combined with AF750 (A-21037, Thermo Fisher Scientific) or QC-1 (custom made by Li-Cor) conjugated to goat anti-mouse secondary antibody at different A:D ratios. While keeping the donor concentration constant at 50µg/mL, the multi-well plate samples contained respective A:D of 0:1, 1:1, 2:1, 3:1, 1:0, 2:0 and 3:0. PBS was used as the solvent and 2 out of the 9 total wells were filled only with PBS to act as negative controls.

The sample was imaged using a single-pixel hyperspectral fluorescence lifetime imaging system developed by Pian et al.^45^ The sample was excited at 695nm using a Mai Tai HP (High-Performance, Mode-Locked, Ti:Sapphire Laser with a repetition rate of 80MHz). The system is composed of structured illumination and detection through a DMD arrangement and is capable of hyperspectrally describing the mentioned sample through the use of PMT-TCSPC based detection. The 16-channel PMT allows for hyperspectral detection ranging from 715 to 780 nm. Each detection channel is approximately 4.5 nm apart of each other. A 715 nm long pass filter (Semrock, FF01-715/LP-25) was used to remove the excitation illumination from the detected signal. The sample was exposed for 1.5s per pattern and a total of 512 frequency arranged Hadamard patterns were acquired to reconstruct 64×64 resolution intensity and lifetime images. Continuous wave (CW) and Time-Domain (TD) reconstructions were retrieved using TVAL3.^57^ The TD reconstructions per detection channel were later used to quantify lifetime using a bi-exponential fitting model that is later detailed throughout this paper.

### Wide-field Macroscopic Fluorescence Lifetime Imaging Platform

MFLI was performed on a wide-field time-domain fluorescence lifetime imaging tomographic system, as described previously.^58^ Briefly, the system excitation source was a tunable Ti-Sapphire laser (Mai Tai HP, Spectra-Physics, CA). The spectral range was 690 – 1040 nm with 100-fs pulses at 80 MHz. The laser was coupled to a digital micro-mirror (DMD) device (DLi 4110, Texas Instruments, TX), which produced a wide-field illumination over an 8×6cm area with 256 grayscale levels encoding at each pixel. The wide-field illumination was spatially modulated by controlling the DMD via Microsoft PowerPoint to ensure optimal signal detection over the whole animal body.^59, 60^ The detection system was a time-gated intensified CCD (ICCD) camera (Picostar HR, Lavision GmbH, Germany). The gate width on the ICCD camera was set to 300 ps. The Instrument Response Function (IRF) and fluorescence signals were collected with 40 ps time interval over a 2.0 ns time window and a 7.0 ns time window, respectively. The total number of gates acquired was 175 and the maximum range of detection was 4096 photons per pixel per gate. The multichannel plate gain (MCP) employed for signal amplification was set to 550 V for the whole imaging session. In this study, imaging was performed in reflectance mode. The laser excitation for AF700 was set at 695 nm and the emission filters were 720±6.5 nm (FF01-720/13-25, Semrock, IL) and 715 nm long pass filter (Semrock, FF01-715/LP-25). The laser for IRDye 800CW was set to 750 nm and the emission filters were 820±6 nm (Semrock, FF01-820/12-25) and 810±45 nm (Chroma Technology, ET810/90). The IRF was measured by using a full field pattern projected on diffuse white paper and acquiring the temporal point spread function (TPSF) without using an emission filter. The imaging platform was equipped with an isoflurane anesthesia machine and a warming device, as described in.^14^

### Microscopy internalization assay of Tf QC-1 vs Tf AF700 in cancer cells

Breast cancer T47D cells were cultured in Dulbecco’s modified Eagle’s medium (Life Technologies) supplemented with 10% fetal calf serum (ATCC), 4 mM L-glutamine (Life Technologies), 10 mM HEPES (Sigma) and Penicillin/Streptomycin (50Units/mL/50µg/mL, Life Technologies) at 37°C and 5% CO2. For the uptake experiment, cells were plated on MatTek 35 mm glass bottom plate (Ashland, MA) 400,000 cells per plate and cultured overnight followed by 30 min incubation with DHB imaging medium (phenol red-free DMEM, 5 mg/mL bovine serum albumin (Sigma), 4 mM L-glutamine, 20 mM HEPES (Sigma) pH 7.4). After that the cells were incubated for 1 h at 37°C with 40 µg/mL either Tf-AF700 or Tf-QC-1 diluted in DHB solution. Tf internalization was stopped by washing the cells with cold HBSS buffer (Life Technologies), followed by fixing for 15 min with 4% paraformaldehyde (PFA) and processing the samples for immunocytochemistry using mouse monoclonal Tf antibody (Serotech 9100-1057) and rabbit polyclonal TfR antibody (Abcam ab84036) diluted 1:250. Secondary antibodies F(ab) goat anti mouse AF555 and goat anti rabbit AF488 (Life Technologies) were used at 1:500 dilution. Samples were imaged on Zeiss LSM 880 confocal microscope using the same settings for all samples. Collected z-stacks were analyzed pixel by pixel for colocalization using Pearson’s coefficient via Imaris software.

### NIR FLIM FRET microscopy

T47D breast cancer cells were plated on MatTek 35 mm glass bottom plates and cultured overnight. Cells were washed with HBSS buffer, precleared for 30 min with DHB imaging medium followed by incubation with Tf FRET pair’s probes for 1 h at 37°C: Tf-AF700 (20 µg/mL), Tf-AF750 (40 ug/mL), Tf-QC-1 (40µg/mL) using Aceptor:Donor ratio 0:1, 0.5:1, 1:1, 2:1 and 3:1. After Tf internalization, cells were washed with cold HBSS buffer, fixed with 4% PFA, and stored in DHB solution for imaging. NIR FLIM FRET microscopy was performed on a Zeiss LSM 880 Airyscan NLO multiphoton confocal microscope (Suppl. Fig.2) using an HPM-100-40 high speed hybrid FLIM detector (GaAs 300-730 nm; Becker & Hickl) and a Titanium: Sapphire laser (Ti: Sa) (680-1040 nm; Chameleon Ultra II, Coherent, Inc.). The Ti: Sa laser was used in conventional one-photon excitation mode. Because of this, the FLIM detector was attached to the confocal output of the scan head. On the LSM 880 with Airyscan, the confocal output from the scan head is used for the ‘Airy-Scan’ detector and thus it is not directly accessible. However, to accommodate the HPM-100-40 detector a Zeiss switching mirror was inserted between the scan head and the Airyscan detector. The 90° position of the switching mirror directs the beam to a vertical port to which the FLIM detector was attached via a Becker & Hickl beamsplitter assembly. A Semrock FF01-716/40 band pass filter and a Semrock FF01-715/LP blocking edge long-pass filter were inserted in the beamsplitter assembly to detect the emission from AF700 and to block scattered light, respectively. The 80/20 beamsplitter in the internal beamsplitter wheel in the LSM 880 was used to direct the 690 nm excitation light to the sample and to pass the emission fluorescence to the FLIM detector. The data was analyzed by two-component exponential fitting using SPCImage software (Becker & Hickl GmbH, Germany). A χ*2* fitness test was used to determine the validity of the fit, providing χ*2* values <1.5 for all pixels.

### Animal experiments

All animal procedures were conducted with the approval of the Institutional Animal Care and Use Committee at both Albany Medical College and Rensselaer Polytechnic Institute. Animal facilities of both institutions have been accredited by the American Association for Accreditation for Laboratory Animals Care International. To produce Matrigel plugs, T47D cells were plated in 60 mm dishes and cultured until confluent. The cells were washed with HBSS buffer and precleared with DHB medium for 30 min. The cells were then loaded with Tf labeled with the appropriate FRET pair AF700/QC or AF700/AF750 (at 40µg/mL constant donor concentration) at A:D ratios of 0:1, 1:1, and 2:1 for 1 h. After internalization, the cells were fixed with 4% PFA, scraped, pelleted, and resuspended in Matrigel (Corning). The 100 µL cell samples in Matrigel (1:1 ratio) were injected into the inguinal and thoracic fat pads on both sides of the anesthetized athymic nude mice for both FRET pairs. Matrigel plugs were allowed to solidify for two hours before imaging.

Tumor xenografts were generated by injecting 5×10^6^ T47D cells or 1×10^6^ MDA-MB 231 cells in phosphate-buffered saline (PBS) mixed 1:1 with Cultrex BME (R&D Systems Inc, Minneapolis, MN, USA) into the right inguinal mammary fat pad of female 5-week old athymic nude mice (NU(NCr)-Foxn1^nu^, Charles River Laboratories, Wilmington, MA, USA). The subcutaneous tumors were allowed to grow for 4-5 weeks and were monitored daily. AF700 and Tf-AF750 (40 and 80 µg respectively in the volume 100-120 µL) were injected through the lateral tail vein in animals restrained with Decapicones.

For IRDye 800CW 2-DG (2-DG; Li-Cor) imaging 100uL of probe (10 nmol/animal) was tail-vein injected in the mice subjected to overnight fasting. In 6 h post injection, the normal feeding was resumed. The animals were imaged 24 h p.i. using MFLI platform. The excitation was set to 750 nm, and the emission filter were 820±6 nm (Semrock, FF01-820/12-25) and 810±45 (Chroma Technology, ET810/90). The imaging parameters were set the same for all mice.

### Immunohistochemistry

After imaging tumors were excised, fixed in formalin, and paraffin embedded. Epitope retrieval was performed by boiling deparaffinized and rehydrated 5µm sections in 10 mM Sodium citrate pH 6.0 for 20 min. IHC staining was carried out using a standard protocol from Vectastain ABC Elite kit (Vector Labs, Burlingame, CA cat#PK-6101). Vector NovaRED (Vector Labs) was used as a peroxidase substrate. Tissue sections were counterstained with Methyl Green (Sigma, cat# M8884). Hematoxylin Eosin stain was used for basic histology. Primary antibodies were as followed: rabbit polyclonal Tf 1:2,000 for IHC (Abcam, cat#1223), rabbit polyclonal TfR 1:250 (Abcam, cat# 84036), rabbit polyclonal GLUT-1 1:200 (ThermoFisher Scientific, cat#PA1-46152). Brightfield images were acquired using Olympus BX40 microscope equipped with Infinity 3 camera (Lumenera Inc., Ottawa, ON, Canada).

### Bi-exponential fitting to extract FRETing donor fraction

In a FRET sample, there are two populations of donor: FRETing and non-FRETing. In order to quantify the amount of FRETing donor within a region of interest, the fluorescence decays in each pixel of a region of interest (ROI) within the sample are analyzed by fitting to a bi-exponential model:

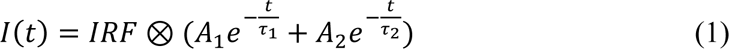

Here I(t) represents the fluorescence decay, IRF is the instrument response function inherent to the system and collected on a piece of diffuse white paper, *A*_1_ and *A*_2_ are the FRETing and non-FRETing donor fractions respectively, τ_1_ and τ_2_ are the quenched and unquenched donor lifetimes, t is time, and ⊗ represents convolution. The tail portion of the decays (99-2% of the peak value) in each pixel of an ROI are fit to the bi-exponential model to extract the FRETing donor fraction (henceforth denoted *A*_1_) using the Matlab function fmincon for least squares minimization of the cost function. The fluorescence decays from the MFLI system and the hyperspectral system were analyzed with the same fitting parameters and smoothing with Anscombe filtering. The average values and standard deviations are reported.

### Statistical analysis

The statistical significance of the data was tested with unpaired Student’s t tests. Differences were considered significant if the p-value was less than 0.05. Error bars indicate standard deviation or 95% confidence interval. For comparison linear regressions, non-parametric Spearman correlation coefficient was utilized via GraphPad software.

## Author’s contribution

M.B. & X.I conceived the original idea. A.R, M.B. & X.I. designed the research study. A.R, J.M & M.B acquired the FLIM data sets. N.S & M.O. acquired and processed the MFLI data sets. N.S, M.O. and A.R. performed the data processing and analyses of results. A.R, M.B & X.I interpreted the results. All authors contributed to the writing of the manuscript. All authors have given approval to the final version of the manuscript.

## Acknowledgments

Authors thank Sez-Jade Chen for collecting antibody binding dataset, Dr. Ling Wang for the help with data analysis and AMC imaging core for the use LSM 880 confocal microscope and help to collect NIR FLIM FRET data. This work was funded by the National Institutes of Health grants R01EB019443, R01CA207725 and R01CA237267.

---

**Supplementary Figure 1.**
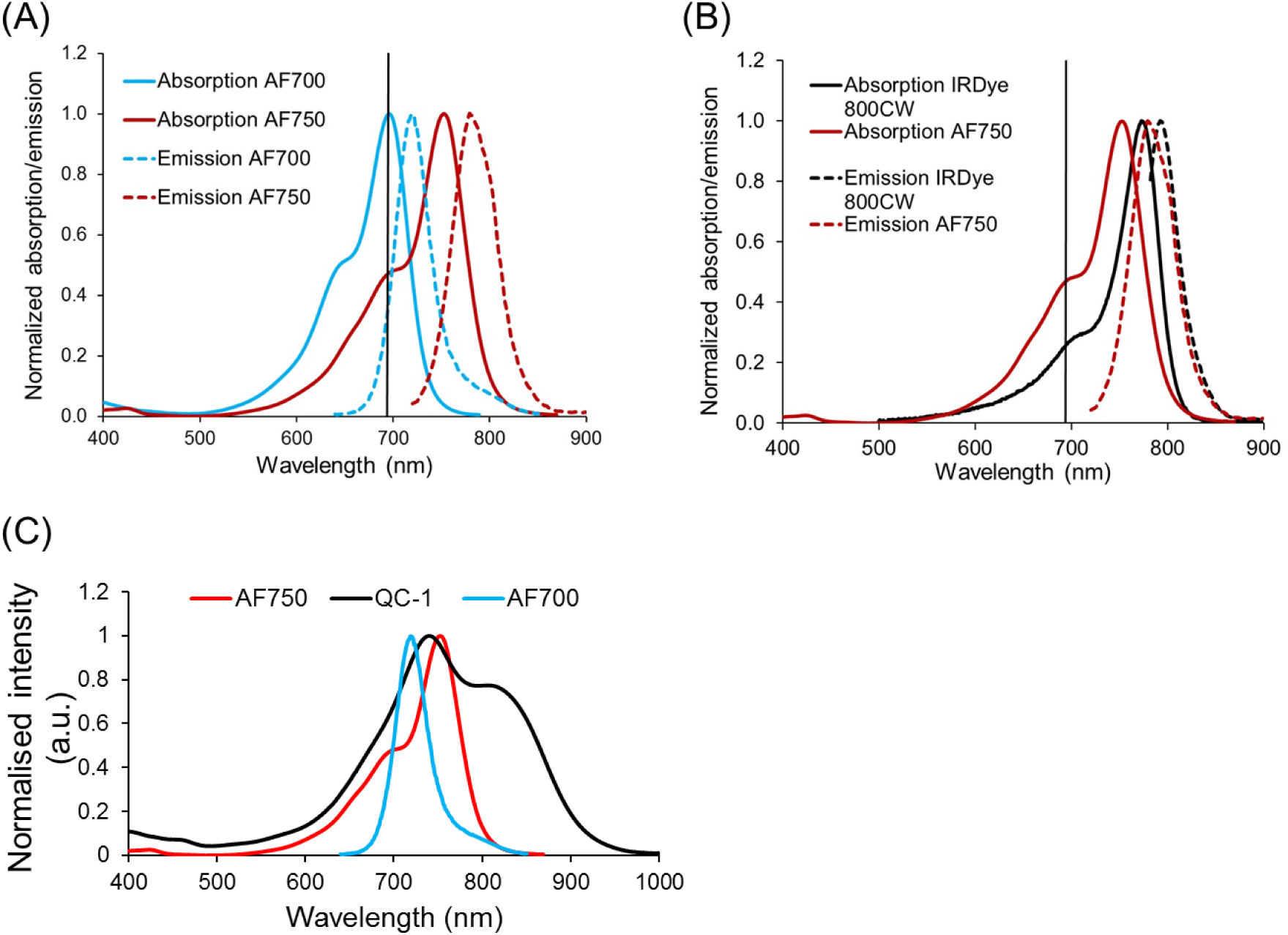
(A) Absorption and emission spectra of FRET pair AF700/750. (B) Absorption and emission spectra of AF750 and IRDye 800CW. Vertical line indicates 695 nm excitation used for MFLI imaging. (C) Absorption and emission spectra of two FRET pairs AF700/750 and AF700/QC-1.

**Supplementary Figure 2.**
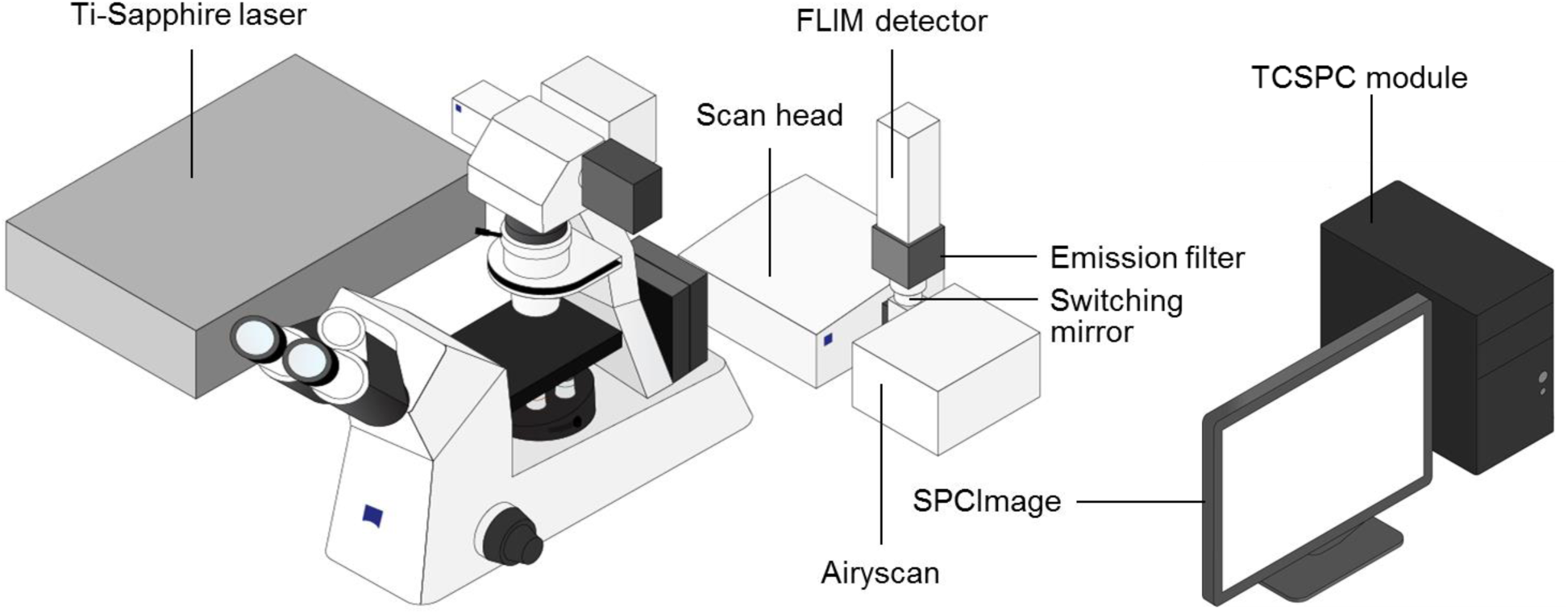
Schematic representation of FLIM system on Zeiss Airyscan multiphoton microscope. Ti: Sapphire laser used in conventional one-photon excitation mode; HPM-100-40 high speed hybrid FLIM detector (Becker & Hickl) is directly coupled to the confocal output of the scan head; a Semrock FF01-716/40 band pass filter and a FF01-715/LP blocking edge long-pass filters are inserted in the beamsplitter assembly to detect the emission from AF700 and to block scattered light, respectively.

**Supplementary Figure 3.**
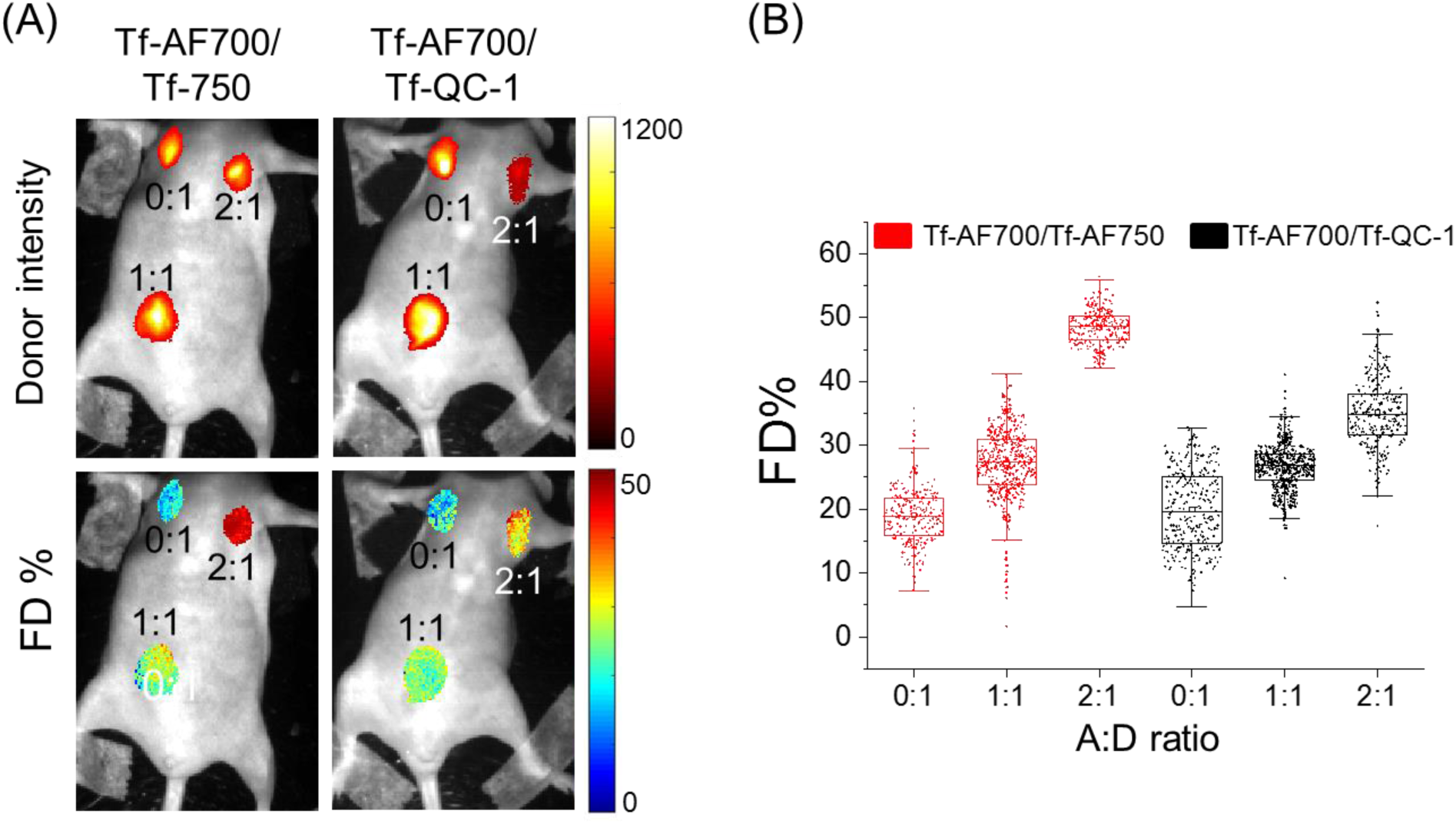
NIR FRET MFLI imaging of Tf-QC-1 as an acceptor in live intact animals. (A) Intensity and FD% maps in Matrigel plugs containing T47D cancer cells preloaded with Tf-AF700/Tf-AF750 or Tf-AF700/Tf-QC-1 ligands with indicated A:D ratio. Anesthetized animals were imaged using wide-field MFLI imager. (B) FRET quantification in cancer cells-Matrigel plugs *in vivo*: box plots showing pixel distribution of FD% at various A:D ratio.

**Supplementary Figure 4.**
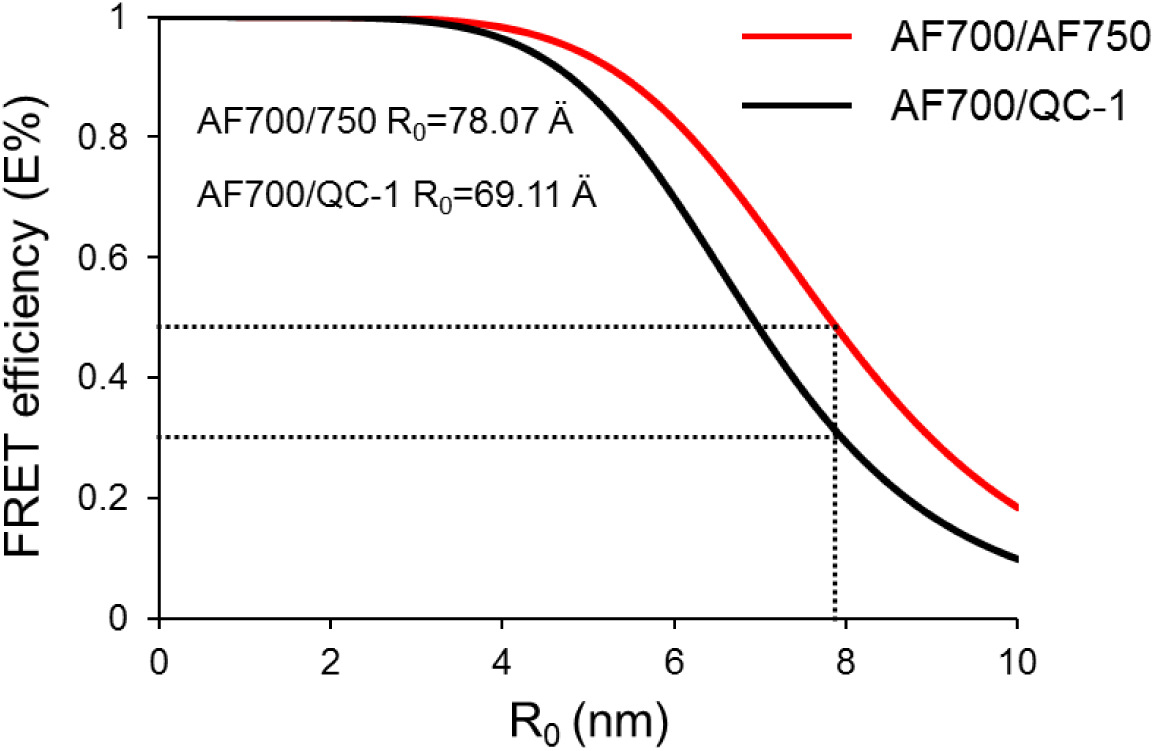
FRET efficiency quantification plot in relation to intrinsic intermolecular distance of AF700/750 and AF700/QC-1.

**Supplementary Figure 5.**
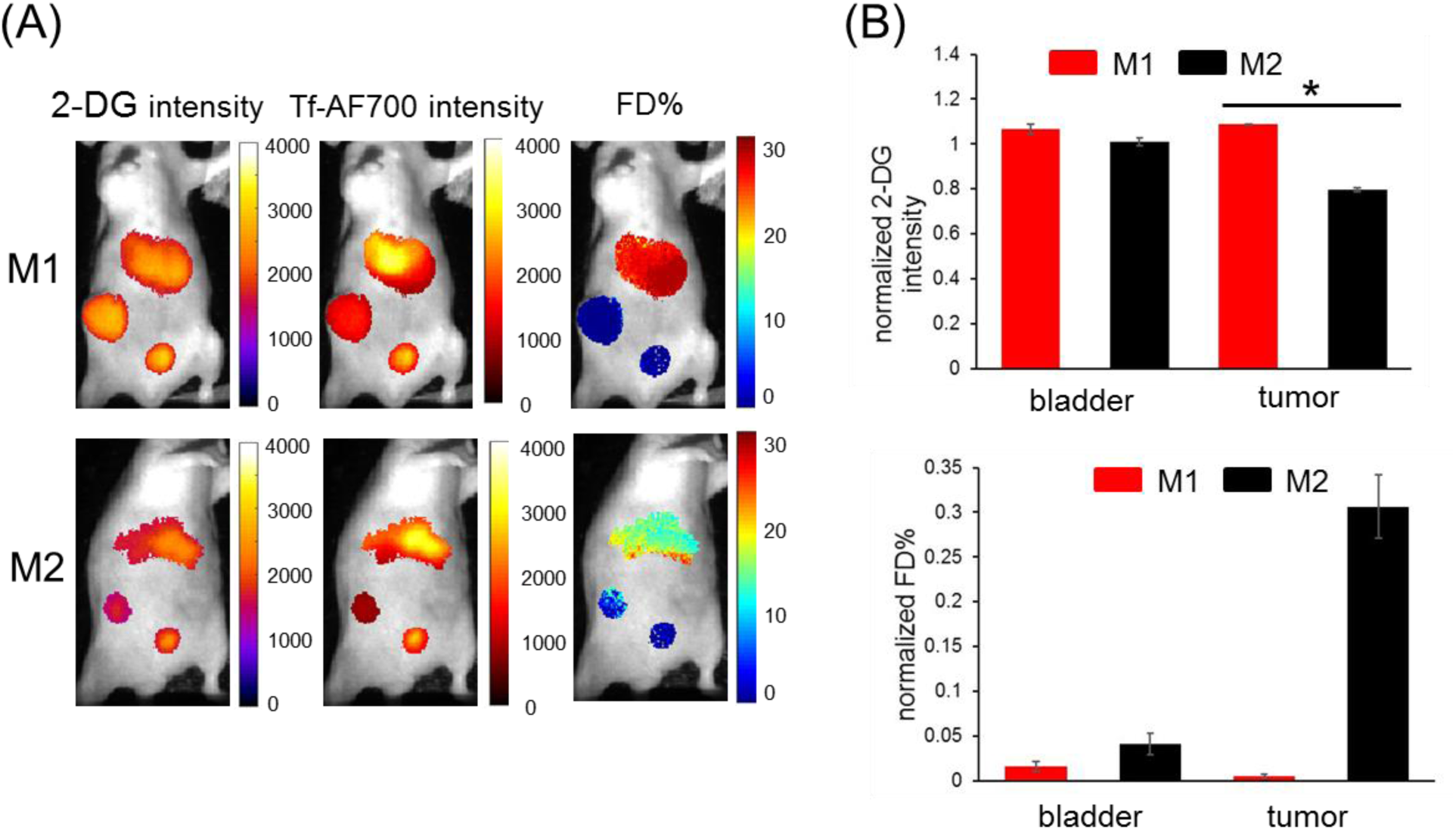
(A) Representative images of 2-DG and Tf-AF700 intensity and FD% map of tumor, liver and urinary bladder in mice simultaneously injected with 2-DG and Tf-AF700/QC-1 and imaged with MFLI FRET at 24 h p.i. (the same animals shown in Fig.7 but with added liver ROIs). (B) Graphs of normalized to liver values 2-DG intensity (upper panel) and FD% (lower panel). Error bars represent 95% confidence interval, asterisks indicate p < 0.05 (p=1.64×10^-42^).

**Supplementary Figure 6.**
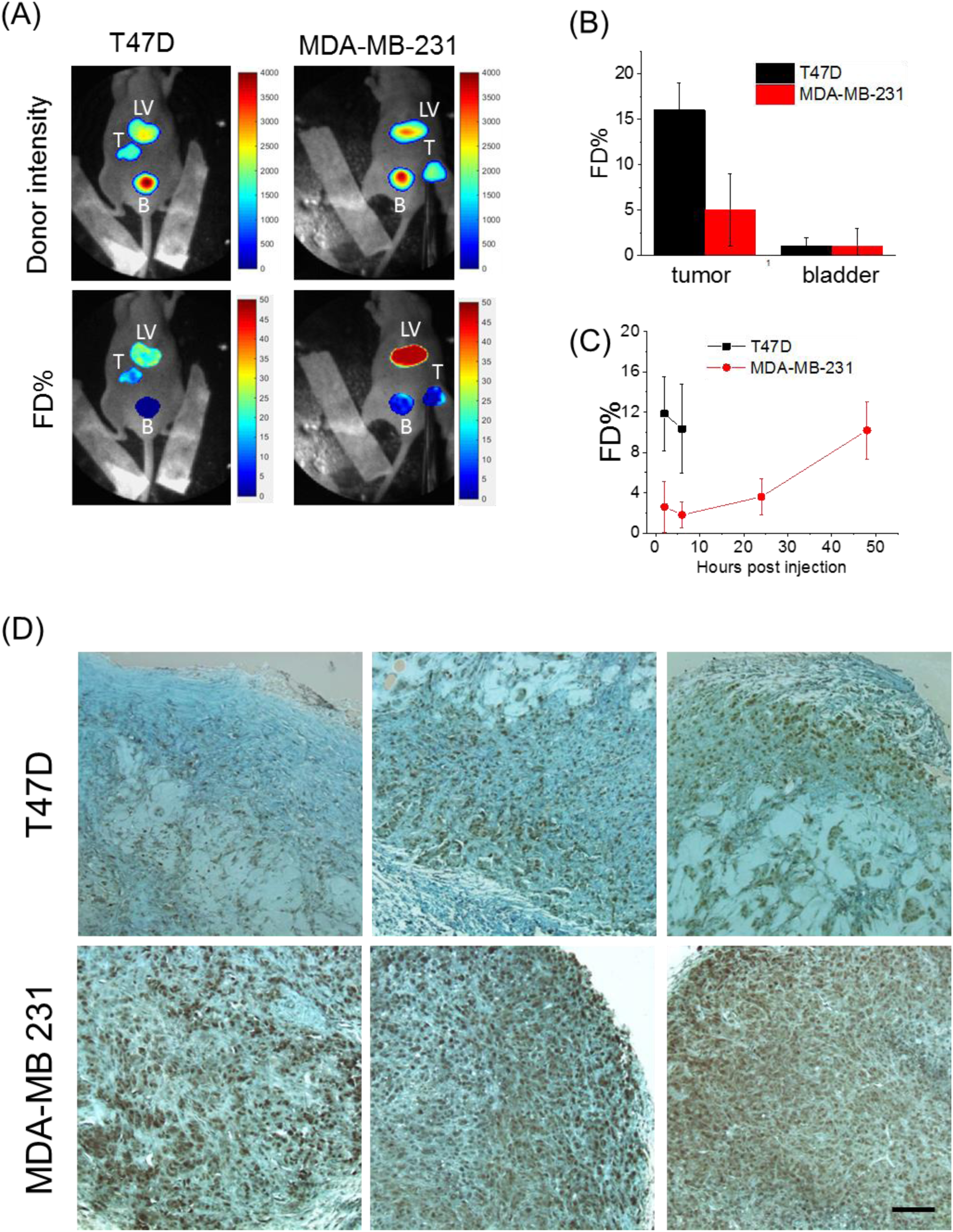
(A) Tumor-carrying mice were injected with 40 µg Tf-AF700 (donor) and 80 µg Tf-AF750 (acceptor) at A:D ratios of 2:1 and subjected to MFLI-FRET imaging at 2 h p.i. Panels show Tf donor fluorescence intensity maximum (total Tf, including soluble and bound Tf) and FD% levels (bound Tf) of live, intact mice using the MFLI-FRET imager. Within each mouse image, ROIs show tumor, liver and bladder fluorescence and FD% levels. (B) Quantification graph FD% in tumor and bladder at 2 h p.i. Error bars indicate standard deviation. (C) Longitudinal imaging of tumors over the course of 48 h MFLI-FRET imaging experiments. N = 5-6 per group from four independent experiment. Data presented as mean + standard deviation. (C) IHC staining of GLUT1 in T47D and MDA-MB-231 tumors derived from 6 different animals. Scale bar 100 µm.

**Supplementary Table 1.**

| Fig. 2 | AF700-QC |  |  |  |  | AF700-AF750 |  |  |  |  |
| --- | --- | --- | --- | --- | --- | --- | --- | --- | --- | --- |
|  | AD 0:1 | AD 0.5:1 | AD 1:1 | AD 2:1 | AD 3:1 | AD 0:1 | AD 0.5:1 | AD 1:1 | AD 2:1 | AD 3:1 |
| Mean FD% | 13.634 | 15.485 | 26.614 | 35.710 | 46.566 | 12.375 | 16.503 | 24.884 | 34.326 | 43.095 |
| Standard deviation | 3.656 | 5.433 | 4.789 | 2.798 | 2.061 | 3.5339 | 4.5035 | 6.265 | 3.129 | 2.576 |
| CI | 0.485 | 0.699 | 0.625 | 0.371 | 0.282 | 0.485 | 0.589 | 0.832 | 0.431 | 0.371 |
| Spearman correlation between two linear regression lines p-value (two-tailed) = 0.0167 |  |  |  |  |  |  |  |  |  |  |

**Supplementary Table 2.**

| Fig. 3 | N | Mean Pearson's coefficient | Standard deviation | SEM | Median |
| --- | --- | --- | --- | --- | --- |
| Tf-AF700 | 5 | 0.6276 | 0.02188 | 0.00978 | 0.6168 |
| Tf-AF750 | 5 | 0.6626 | 0.02146 | 0.0096 | 0.6592 |
| Two tailed t-test p-value 0.0888 |  |  |  |  |  |

**Supplementary Table 3.**

| Fig. 4 | Tf-AF700/Tf-QC1 |  |  |  |  | Tf-AF700/Tf-AF750 |  |  |  |  |
| --- | --- | --- | --- | --- | --- | --- | --- | --- | --- | --- |
| A:D ratio | 0:1 | 0.5:1 | 1:1 | 2:1 | 3:1 | 0:1 | 0.5:1 | 1:1 | 2:1 | 3:1 |
| Mean FD% | 23.152 | 40.061 | 44.596 | 52.497 | 54.044 | 23.36 | 46.754 | 55.978 | 64.529 | 68.301 |
| Standard deviation | 9.823 | 3.947 | 3.252 | 3.049 | 6.284 | 8.896 | 4.843 | 5.228 | 6.254 | 4.383 |
| CI | 0.195 | 0.078 | 0.065 | 0.060 | 0.125 | 0.176 | 0.096 | 0.104 | 0.124 | 0.087 |
| Two- sample t-test p-value |  |  |  |  |  | 0.96 | 0.0034 | 1.5x10<sup>-5</sup> | 3.4x10<sup>-5</sup> | 1.4x10<sup>-5</sup> |

**Supplementary Table 4.**

| Fig.7<br>tumors | N total | Mean | Standard<br>Deviation | tumors | N total | Mean | Standard<br>Deviation |
| --- | --- | --- | --- | --- | --- | --- | --- |
| M1 2-DG | 655 | 2308.858 | 354.465 | M1 FD% | 431 | 0.221 | 0.989 |
| M2 2-DG | 250 | 1526.564 | 126.486 | M2 FD% | 244 | 7.936 | 7.293 |
| Two-sample t-test (Welch Correction) p-value<br>1.061x10<sup>-255</sup> |  |  |  | Two-sample t-test (Welch Correction) p-value<br>1.418x10<sup>-41</sup> |  |  |  |

## References

1. O’Farrell, A. C.; Shnyder, S. D.; Marston, G.; Coletta, P. L.; Gill, J. H. Non-Invasive Molecular Imaging for Preclinical Cancer Therapeutic Development. Br. J. Pharmacol. 2013, 169 (4), 719–735. https://doi.org/10.1111/bph.12155.

2. Waaijer, S. J. H.; Kok, I. C.; Eisses, B.; Schröder, C. P.; Jalving, M.; Brouwers, A. H.; Lub-de Hooge, M. N.; de Vries, E. G. E. Molecular Imaging in Cancer Drug Development. J. Nucl. Med. 2018, 59 (5), jnumed.116.188045. https://doi.org/10.2967/jnumed.116.188045.

3. Lindner, J. R.; Link, J. Molecular Imaging in Drug Discovery and Development. Circ. Cardiovasc. Imaging 2018, 11 (2), 1–11. https://doi.org/10.1161/CIRCIMAGING.117.005355.

4. Broos, K.; Lecocq, Q.; Raes, G.; Devoogdt, N.; Keyaerts, M.; Breckpot, K. Noninvasive Imaging of the PD-1: PD-L1 Immune Checkpoint: Embracing Nuclear Medicine for the Benefit of Personalized Immunotherapy. Theranostics 2018, 8 (13), 3559–3570. https://doi.org/10.7150/thno.24762.

5. Truillet, C.; Oh, H. L. J.; Yeo, S. P.; Lee, C. Y.; Huynh, L. T.; Wei, J.; Parker, M. F. L.; Blakely, C.; Sevillano, N.; Wang, Y. H.;, et al. Imaging PD-L1 Expression with ImmunoPET. Bioconjug. Chem. 2018. https://doi.org/10.1021/acs.bioconjchem.7b00631.

6. Kumar, D.; Lisok, A.; Dahmane, E.; McCoy, M.; Shelake, S.; Chatterjee, S.; Allaj, V.; Sysa-Shah, P.; Wharram, B.; Lesniak, W. G.;, et al. Peptide-Based PET Quantifies Target Engagement of PD-L1 Therapeutics. J. Clin. Invest. 2019. https://doi.org/10.1172/JCI122216.

7. Lee, S. S.-Y.; Bindokas, V. P.; Kron, S. J. Multiplex Three-Dimensional Mapping of Macromolecular Drug Distribution in the Tumor Microenvironment. Mol. Cancer Ther. 2018. https://doi.org/10.1158/1535-7163.mct-18-0554.

8. Burvenich, I. J. G.; Parakh, S.; Parslow, A. C.; Lee, S. T.; Gan, H. K.; Scott, A. M. Receptor Occupancy Imaging Studies in Oncology Drug Development. AAPS J. 2018, 20 (2), 43. https://doi.org/10.1208/s12248-018-0203-z.

9. Durham, T. B.; Blanco, M. J. Target Engagement in Lead Generation. Bioorganic and Medicinal Chemistry Letters. 2015. https://doi.org/10.1016/j.bmcl.2014.12.076.

10. Molina, D. M.; Jafari, R.; Ignatushchenko, M.; Seki, T.; Larsson, E. A.; Dan, C.; Sreekumar, L.; Cao, Y.; Nordlund, P. Monitoring Drug Target Engagement in Cells and Tissues Using the Cellular Thermal Shift Assay. Science (80-.). 2013. https://doi.org/10.1126/science.1233606.

11. Dubach, J. M.; Kim, E.; Yang, K.; Cuccarese, M.; Giedt, R. J.; Meimetis, L. G.; Vinegoni, C.; Weissleder, R. Quantitating Drug-Target Engagement in Single Cells in Vitro and in Vivo. Nat. Chem. Biol. 2017, 13 (2), 168–173. https://doi.org/10.1038/nchembio.2248.

12. Berezin, M. Y.; Achilefu, S. Fluorescence Lifetime Measurements and Biological Imaging. Chem. Rev. 2010. https://doi.org/10.1021/cr900343z.

13. Rajoria, S.; Zhao, L.; Intes, X.; Barroso, M. FLIM-FRET for Cancer Applications. Curr. Mol. Imaging 2015, 3 (2), 144–161. https://doi.org/10.2174/2211555203666141117221111.

14. Abe, K.; Zhao, L.; Periasamy, A.; Intes, X.; Barroso, M. Non-Invasive in Vivo Imaging of near Infrared-Labeled Transferrin in Breast Cancer Cells and Tumors Using Fluorescence Lifetime FRET. PLoS One 2013, 8 (11). https://doi.org/10.1371/journal.pone.0080269.

15. Wallrabe, H.; Elangovan, M.; Burchard, A.; Periasamy, A.; Barroso, M. Confocal FRET Microscopy to Measure Clustering of Ligand-Receptor Complexes in Endocytic Membranes. Biophys. J. 2003, 85 (1), 559–571. https://doi.org/10.1016/S0006-3495(03)74500-7.

16. Wallrabe, H.; Periasamy, A.; Kim, C.; Talati, R.; Barroso, M. Confocal FRET and FLIM Microscopy to Characterize the Distribution of Membrane Receptors. Proc. SPIE 2006, 6089, 608905.

17. Wallrabe, H.; Bonamy, G.; Periasamy, A.; Barroso, M. Receptor Complexes Cotransported via Polarized Endocytic Pathways Form Clusters with Distinct Organizations. Mol. Biol. Cell 2007. https://doi.org/10.1091/mbc.e06-08-0700.

18. Talati, R.; Vanderpoel, A.; Eladdadi, A.; Anderson, K.; Abe, K.; Barroso, M. Automated Selection of Regions of Interest for Intensity-Based FRET Analysis of Transferrin Endocytic Trafficking in Normal vs. Cancer Cells. Methods 2014, 66 (2), 139–152. https://doi.org/10.1016/j.ymeth.2013.08.017.

19. Mazurkiewicz, J. E.; Herrick-Davis, K.; Barroso, M.; Ulloa-Aguirre, A.; Lindau-Shepard, B.; Thomas, R. M.; Dias, J. A. Single-Molecule Analyses of Fully Functional Fluorescent Protein-Tagged Follitropin Receptor Reveal Homodimerization and Specific Heterodimerization with Lutropin Receptor1. Biol. Reprod. 2015. https://doi.org/10.1095/biolreprod.114.125781.

20. Barroso, M.; Tucker, H.; Drake, L.; Nichol, K.; Drake, J. R. Antigen-B Cell Receptor Complexes Associate with Intracellular Major Histocompatibility Complex (MHC) Class II Molecules. J. Biol. Chem. 2015, 290 (45), 27101–27112. https://doi.org/10.1074/jbc.M115.649582.

21. Elangovan, M.; Wallrabe, H.; Chen, Y.; Day, R. N.; Barroso, M.; Periasamy, A. Characterization of One- and Two-Photon Excitation Fluorescence Resonance Energy Transfer Microscopy. Methods 2003. https://doi.org/10.1016/S1046-2023(02)00283-9.

22. Rudkouskaya, A.; Sinsuebphon, N.; Ward, J.; Tubbesing, K.; Intes, X.; Barroso, M. Quantitative Imaging of Receptor-Ligand Engagement in Intact Live Animals. J. Control. Release 2018, 286, 451–459. https://doi.org/10.1016/j.jconrel.2018.07.032.

23. Chen, S. J.; Sinsuebphon, N.; Rudkouskaya, A.; Barroso, M.; Intes, X.; Michalet, X. In Vitro and in Vivo Phasor Analysis of Stoichiometry and Pharmacokinetics Using Short-Lifetime near-Infrared Dyes and Time-Gated Imaging. J. Biophotonics 2019. https://doi.org/10.1002/jbio.201800185.

24. Sinsuebphon, N.; Rudkouskaya, A.; Barroso, M.; Intes, X. Comparison of Illumination Geometry for Lifetime-Based Measurements in Whole-Body Preclinical Imaging. J. Biophotonics 2018, No. May, jbio.201800037. https://doi.org/10.1002/jbio.201800037.

25. Kovar, J. L.; Volcheck, W.; Sevick-Muraca, E.; Simpson, M. A.; Olive, D. M. Characterization and Performance of a Near-Infrared 2-Deoxyglucose Optical Imaging Agent for Mouse Cancer Models. Anal. Biochem. 2009. https://doi.org/10.1016/j.ab.2008.09.050.

26. Voigt, W. Advanced PET Imaging in Oncology: Status and Developments with Current and Future Relevance to Lung Cancer Care. Current Opinion in Oncology. 2018. https://doi.org/10.1097/CCO.0000000000000430.

27. Shcherbakova, D. M.; Cox Cammer, N.; Huisman, T. M.; Verkhusha, V. V.; Hodgson, L. Direct Multiplex Imaging and Optogenetics of Rho GTPases Enabled by Near-Infrared FRET Article. Nat. Chem. Biol. 2018. https://doi.org/10.1038/s41589-018-0044-1.

28. Hoffmann, K.; Behnke, T.; Drescher, D.; Kneipp, J.; Resch-Genger, U. Near-Infrared-Emitting Nanoparticles for Lifetime-Based Multiplexed Analysis and Imaging of Living Cells. ACS Nano 2013. https://doi.org/10.1021/nn4029458.

29. Grant, D. M.; Zhang, W.; McGhee, E. J.; Bunney, T. D.; Talbot, C. B.; Kumar, S.; Munro, I.; Dunsby, C.; Neil, M. A. A.; Katan, M.;, et al. Multiplexed FRET to Image Multiple Signaling Events in Live Cells. Biophys. J. 2008, 95 (10), L69–L71. https://doi.org/10.1529/biophysj.108.139204.

30. Welch, C. M.; Elliott, H.; Danuser, G.; Hahn, K. M. Imaging the Coordination of Multiple Signalling Activities in Living Cells. Nat. Rev. Mol. Cell Biol. 2011, 12 (11), 749–756. https://doi.org/10.1038/nrm3212.

31. Geißler, D.; Stufler, S.; Löhmannsröben, H. G.; Hildebrandt, N. Six-Color Time-Resolved Förster Resonance Energy Transfer for Ultrasensitive Multiplexed Biosensing. J. Am. Chem. Soc. 2013, 135 (3), 1102–1109. https://doi.org/10.1021/ja310317n.

32. Busch, C.; Schroter, T.; Grabolle, M.; Wenzel, M.; Kempe, H.; Kaiser, W. A.; Resch-Genger, U.; Hilger, I. An In Vivo Spectral Multiplexing Approach for the Cooperative Imaging of Different Disease-Related Biomarkers with Near-Infrared Fluorescent Forster Resonance Energy Transfer Probes. J. Nucl. Med. 2012. https://doi.org/10.2967/jnumed.111.094391.

33. Le Reste, L.; Hohlbein, J.; Gryte, K.; Kapanidis, A. N. Characterization of Dark Quencher Chromophores as Nonfluorescent Acceptors for Single-Molecule FRET. Biophys. J. 2012, 102 (11), 2658–2668. https://doi.org/10.1016/j.bpj.2012.04.028.

34. Myochin, T.; Hanaoka, K.; Iwaki, S.; Ueno, T.; Komatsu, T.; Terai, T.; Nagano, T.; Urano, Y. Development of a Series of Near-Infrared Dark Quenchers Based on Si-Rhodamines and Their Application to Fluorescent Probes. J. Am. Chem. Soc. 2015, 137 (14), 4759–4765. https://doi.org/10.1021/jacs.5b00246.

35. Johansson, M. K. Choosing Reporter-Quencher Pairs for Efficient Quenching through Formation of Intramolecular Dimers. Methods Mol. Biol. 2006, 335, 17–29. https://doi.org/10.1385/1-59745-069-3:17.

36. Peng, X.; Chen, H.; Draney, D. R.; Volcheck, W.; Schutz-Geschwender, A.; Olive, D. M. A Nonfluorescent, Broad-Range Quencher Dye for Förster Resonance Energy Transfer Assays. Anal. Biochem. 2009, 388 (2), 220–228. https://doi.org/10.1016/j.ab.2009.02.024.

37. Osterman, H. The Next Step in Near Infrared Fluorescence: IRDye QC-1 Dark Quencher. Rev. Artic. 2009, 388 (2009), 1–8.

38. Gong, H.; Urlacher, T. A Homogeneous Fluorescence-Based Method to Measure Antibody Internalization in Tumor Cells. Anal. Biochem. 2015, 469, 1–3. https://doi.org/10.1016/j.ab.2014.09.008.

39. Obaid, G.; Spring, B. Q.; Bano, S.; Hasan, T. Activatable Clinical Fluorophore-Quencher Antibody Pairs as Dual Molecular Probes for the Enhanced Specificity of Image-Guided Surgery. J. Biomed. Opt. 2017, 22 (12), 1. https://doi.org/10.1117/1.JBO.22.12.121607.

40. Yim, J. J.; Tholen, M.; Klaassen, A.; Sorger, J.; Bogyo, M. Optimization of a Protease Activated Probe for Optical Surgical Navigation. Mol. Pharm. 2017. https://doi.org/10.1021/acs.molpharmaceut.7b00822.

41. Zhang, J.; Smaga, L. P.; Satyavolu, N. S. R.; Chan, J.; Lu, Y. DNA Aptamer-Based Activatable Probes for Photoacoustic Imaging in Living Mice. J. Am. Chem. Soc. 2017, 139 (48), 17225–17228. https://doi.org/10.1021/jacs.7b07913.

42. Joseph, J.; Koehler, P.; Zuehlsdorff, T. J.; Cole, D. J.; Baumann, K. N.; Weber, J.; Bohndiek, S. E.; Hernández-ainsa, S. Photoacoustic Molecular Rulers Based on DNA Nanostructures. 2017, 1–15. https://doi.org/10.1101/125583.

43. Qian, Z. M. M.; Li, H.; Sun, H.; Ho, K. Targeted Drug Delivery via the Transferrin Receptor-Mediated Endocytosis Pathway. Pharmacol. Rev. 2002, 54 (4), 561–587. https://doi.org/10.1124/pr.54.4.561.

44. Daniels, T. R.; Delgado, T.; Helguera, G.; Penichet, M. L. The Transferrin Receptor Part II: Targeted Delivery of Therapeutic Agents into Cancer Cells. Clinical Immunology. 2006. https://doi.org/10.1016/j.clim.2006.06.006.

45. Pian, Q.; Yao, R.; Sinsuebphon, N.; Intes, X. Compressive Hyperspectral Time-Resolved Wide-Field Fluorescence Lifetime Imaging. Nat. Photonics 2017, 11 (7), 411–414. https://doi.org/10.1038/nphoton.2017.82.

46. Holliday, D. L.; Speirs, V. Choosing the Right Cell Line for Breast Cancer Research. Breast Cancer Research. 2011. https://doi.org/10.1186/bcr2889.

47. Mayle, K. M.; Le, A. M.; Kamei, D. T. The Intracellular Trafficking Pathway of Transferrin. Biochimica et Biophysica Acta - General Subjects. 2012. https://doi.org/10.1016/j.bbagen.2011.09.009.

48. Barroso, M.; Sun, Y.; Wallrabe, H.; Periasamy, A. Nanometer-Scale Measurements Using FRET and FLIM Microscopy. In Luminescence; 2013. https://doi.org/10.1201/b15490-11.

49. Szablewski, L. Expression of Glucose Transporters in Cancers. Biochimica et Biophysica Acta - Reviews on Cancer. 2013. https://doi.org/10.1016/j.bbcan.2012.12.004.

50. Carvalho, K. C.; Cunha, I. W.; Rocha, R. M.; Ayala, F. R.; Cajaíba, M. M.; Begnami, M. D.; Vilela, R. S.; Paiva, G. R.; Andrade, R. G.; Soares, F. A. GLUT1 Expression in Malignant Tumors and Its Use as an Immunodiagnostic Marker. Clinics 2011. https://doi.org/10.1590/S1807-59322011000600008.

51. J.; Ye, C.; Chen, C.; Xiong, H.; Xie, B.; Zhou, J.; Chen, Y.; Zheng, S.; Wang, L. Glucose Transporter GLUT1 Expression and Clinical Outcome in Solid Tumors: A Systematic Review and Meta-Analysis. Oncotarget 2017. https://doi.org/10.18632/oncotarget.15171.

52. Airley, R.; Loncaster, J.; Bromley, M.; Roberts, S.; West, C.; Davidson, S.; Hunter, R.; Airley, R.; Patterson, A.; Stratford, I.;, et al. Glucose Transporter Glut-1 Expression Correlates with Tumor Hypoxia and Predicts Metastasis-Free Survival in Advanced Carcinoma of the Cervix. Clin. Cancer Res. 2001.

53. Grantab, R. H.; Tannock, I. F. Penetration of Anticancer Drugs through Tumour Tissue as a Function of Cellular Packing Density and Interstitial Fluid Pressure and Its Modification by Bortezomib. BMC Cancer 2012. https://doi.org/10.1186/1471-2407-12-214.

54. Kwon, I. K.; Lee, S. C.; Han, B.; Park, K. Analysis on the Current Status of Targeted Drug Delivery to Tumors. J. Control. Release 2012, 164 (2), 108–114. https://doi.org/10.1016/j.jconrel.2012.07.010.

55. Kim, M.; Kim, D.; Jung, W.; Koo, J. Expression of Metabolism-Related Proteins in Triple-Negative Breast Cancer. 2014, 7 (1), 301–312.

56. Kaira, K.; Higuchi, T.; Naruse, I.; Arisaka, Y.; Tokue, A.; Altan, B.; Suda, S.; Mogi, A.; Shimizu, K.; Sunaga, N.;, et al. Metabolic Activity By18F–FDG-PET/CT Is Predictive of Early Response after Nivolumab in Previously Treated NSCLC. Eur. J. Nucl. Med. Mol. Imaging 2018, 45 (1), 56–66. https://doi.org/10.1007/s00259-017-3806-1.

57. Li, C. Compressive Sensing for 3D Data Processing Tasks : Applications, Models and Algorithms. Rice Univ. 2011.

58. Venugopal, V.; Chen, J.; Barroso, M.; Intes, X. Quantitative Tomographic Imaging of Intermolecular FRET in Small Animals. Biomed. Opt. Express 2012, 3 (12), 3161–3175. https://doi.org/10.1364/BOE.3.003161.

59. Zhao, L.; Abe, K.; Rajoria, S.; Pian, Q.; Barroso, M.; Intes, X. Spatial Light Modulator Based Active Wide-Field Illumination for Ex Vivo and in Vivo Quantitative NIR FRET Imaging. Biomed. Opt. Express 2014, 5 (3), 944. https://doi.org/10.1364/BOE.5.000944.

60. Zhao, L.; Abe, K.; Barroso, M.; Intes, X. Active Wide-Field Illumination for High-Throughput Fluorescence Lifetime Imaging. Opt. Lett. 2013, 38 (19), 3976. https://doi.org/10.1364/OL.38.003976.

